# Structure of chlorophyll synthase in complex with the LHC-like protein HliD

**DOI:** 10.64898/2026.05.13.724888

**Authors:** Dmitry Shvarev, Anna Wysocka, Felix S. Morey-Burrows, Karolina O. Panas, Arman Pazuki, Natalia Kulik, Matthew S. Proctor, Jan Pilný, C. Neil Hunter, Andrew Hitchcock, Roman Sobotka

**Affiliations:** Department of Biology/Chemistry - Section of Structural Biology, University of Osnabrueck, Osnabrueck 49076, Germany; Department of Biology/Chemistry - Section of Structural Biology of Photosynthetic Microorganisms, University of Osnabrueck, Osnabrueck 49076, Germany; Center of Cellular Nanoanalytic Osnabrueck (CellNanOs); Osnabrueck University, Osnabrueck 49076, Germany; Centre Algatech, Institute of Microbiology of the Czech Academy of Sciences, Třeboň 37901, Czech Republic; Faculty of Science, University of South Bohemia, České Budějovice, 370 05, Czech Republic; Plants, Photosynthesis and Soil, School of Biosciences, University of Sheffield, Sheffield S10 2TN, United Kingdom; Molecular Microbiology: Biochemistry to Disease, School of Biosciences, University of Sheffield, Sheffield S10 2TN, United Kingdom

## Abstract

Chlorophyll is the central cofactor in oxygenic photosynthesis, responsible for both light capture in antenna complexes and light-driven charge separation in photosystems. The tetraprenyl tail of chlorophyll is attached to the chlorophyllide macrocycle by the transmembrane enzyme chlorophyll synthase (ChlG). In the cyanobacterium *Synechocystis* sp. PCC 6803, ChlG forms a complex with the LHC-like high-light-inducible protein HliD. To understand the substrate specificity and catalytic mechanism of ChlG, and how it is photoprotected by HliD, we determined the structure of a ChlG_2_HliD_2_ complex in both substrate-free and geranylgeranyl pyrophosphate (GGPP)-bound states using cryogenic electron microscopy (cryo-EM). A homodimer of HliD, which binds four chlorophylls and two zeaxanthin molecules in a quenched state, is flanked by two ChlG monomers. AlphaFold modelling placed chlorophyllide adjacent to the structurally resolved GGPP bound to the active site of ChlG. Cryo-EM data, site-directed mutagenesis and molecular dynamics were used to formulate a molecular mechanism for ChlG catalysis. The structure of the ChlG_2_HliD_2_ complex shows how Hlips bind to the synthase, reveals the arrangements of carotenoids and chlorophylls that mediate energy dissipation, and sheds light on the evolution of eukaryotic LHC antennae from their cyanobacterial ancestors.

## Introduction

Chlorophyll *a* (Chl) (**Extended Data Figure 1a**) is the major light-absorbing pigment on Earth, and it is responsible for funneling solar energy into the biosphere. The universal tetrapyrrole precursor 5-aminolevulinic acid initiates the synthesis of Chl in plants, algae and cyanobacteria ^1^, and the protoporphyrin IX intermediate is the substrate for either chelation of Fe^2+^ leading to formation of heme and bilins, or insertion of Mg^2+^, the first dedicated step in Chl biosynthesis ^1^. Following modifications to the macrocycle and the formation of the isocyclic E ring, Chl biosynthesis is completed by the addition of a polyisoprene alcohol to chlorophyllide *a* (Chlide) by the combined activities of the Chl synthase (ChlG) and geranylgeranyl (GG) reductase (ChlP) enzymes (**Extended Data Figure 1b**) ^2^. ChlG binds the GG pyrophosphate (GGPP) and Chlide substrates and catalyses the esterification of the C17-propionate side chain on ring D by GG. ChlP subsequently catalyses the stepwise reduction of three double bonds in the GG moiety of GG-Chl to phytyl, yielding Chl as the final product ^2^.

The *in vitro* activity of ChlG was first demonstrated in a plant extract ^3^, followed by genetic studies of *bchG*, encoding the equivalent bacteriochlorophyll (BChl) synthase, in the purple phototrophic bacteria *Rhodobacter sphaeroides* and *Rhodobacter capsulatus* ^4,5^. Definite assignments of function were possible by measuring BchG activities in recombinant proteins ^6^. Based on its homology to *bchG*, *chlG* was identified in the sequenced genomes of the model cyanobacterium *Synechocystis* sp. PCC 6803 (hereafter *Synechocystis*) ^7^ and the model flowering plant *Arabidopsis thaliana* (*Arabidopsis*) ^8^; heterologous expression of *Synechocystis chlG* in *Escherichia coli* yielded assayable activities of ChlG ^6^. The synthases exhibit substrate specificities, so BchG uses bacteriochlorophyllide (BChlide) and is inhibited by Chlide, whereas cyanobacterial ChlG is unable to esterify BChlide ^9^. Homologies between BChl and Chl synthases, and their relationship to other polyprenyltransferases, which catalyze the attachment of phytyl or isoprenyl chains to an aromatic ring ^10,11^, were noted early on ^12^. While several structures of the prenyltransferase UbiA, essential for ubiquinone biosynthesis, have been described ^10,11^, no structural information has been available for BchG or ChlG.

In cyanobacteria, ChlG is an essential, intrinsic membrane protein with a predicted molecular mass of approximately 35 kDa that has been isolated as a larger, enzymatically active complex with two high-light-inducible proteins (Hlips), HliC and HliD ^13,14^. Previous studies also showed that ChlG can participate in larger assemblies involving the Alb3/YidC insertase and the Ycf39 protein, a putative assembly factor for photosystem II (PSII) ^13^. Hlips are single transmembrane helix (TMH) proteins that can form homodimers or heterodimers and are predicted to bind four Chl and two carotenoid molecules per dimer ^15^. Based on sequence homology, cyanobacterial Hlips have been proposed as the ancestor of the light-harvesting complexes (LHC) of algae and plants ^16,17^. However, in contrast to LHCs, Hlip-associated pigments are organized in a stable, quenched configuration, with virtually all the energy absorbed by the Chls quickly dissipated by the carotenoids ^18,19^. Although free Hlips exclusively bind β-carotene (β-Car) ^18,20^, isolated ChlG-Hlip complexes contain also zeaxanthin (Zea) and myxoxanthophyll (Myx) ^13^, with the Zea molecules implicated in a structural role stabilizing the ChlG–HliD interaction ^21^.

We showed recently ^22^ that under low stress conditions *Synechocystis* ChlG and Hlips are organized as a hetero-tetrameric complex with two copies of ChlG attached to an HliD dimer (ChlG-HliD_2_-ChlG complex, hereafter G-D_2_-G). Stress conditions (e.g. high light) result in a massive accumulation of HliC, followed by the rapid replacement of G-D_2_-G complexes by trimeric ChlG-HliD-HliC assemblies, which are proposed to play a specific role in the repair of PSII ^22^. It is not known how this potential Chl-delivering machinery works, how it binds to Hlips, or how it engages with the wider assembly apparatus comprising the translocase and YidC insertase ^13^. ChlG must also participate in the repair of PSII ^23^, so structural and mechanistic information is essential to understand its multiple roles.

Here, we present the structures of the G-D_2_-G complex from *Synechocystis* solved by single particle cryogenic electron microscopy (cryo-EM) in both apo (3.0 Å) and GGPP-bound (3.2 Å) forms. The complex contains four Chls, two Zea and two Myx pigments, and, in the case of the GGPP-bound complex, a molecule of GGPP is bound within the active site of each ChlG monomer. We combine the cryo-EM data with structural modelling, molecular dynamics (MD) and functional analyses to propose a catalytic mechanism for esterification of the C17-propionate of Chlide, and our models account for the specificity of ChlG and BchG for their respective Chlide and BChlide substrates. The inspection of HliD-associated pigments reveals a remarkably close similarity with the LHC antennae of plants and algae, consistent with their proposed evolutionary relationship.

## Results

### Overall architecture of the G-D_2_-G complex

The FLAG-tagged G-D_2_-G complex was purified using the protocol described previously ^22,24^. The analysis of the obtained eluate by 2D clear-native (CN)/SDS gel electrophoresis confirmed that the G-D_2_-G complex is highly pure, although it contains a fraction of the smaller G-D_2_ complex corresponding to a monomer of ChlG bound to two copies of HliD (**Extended Data Figure 2**). To determine the structure with the bound substrate, the isolated complex was additionally incubated with 150 µM GGPP in the purification buffer prior to preparation of cryo-EM grids. The structures of the G-D_2_-G complex in substrate-free (apo) and GGPP-bound states were refined to 3.0-3.4 Å and 3.2-3.6 Å, respectively (**Figure 1, Extended Data Figures 3-6, Table S1**). In both cases, the extended dimeric complex, 124-126 Å in length, displays almost identical organization and contains two ChlG monomers on the periphery linked through two crossed HliD monomers at the center of the structure (**Figure 1**).

**Figure 1.**
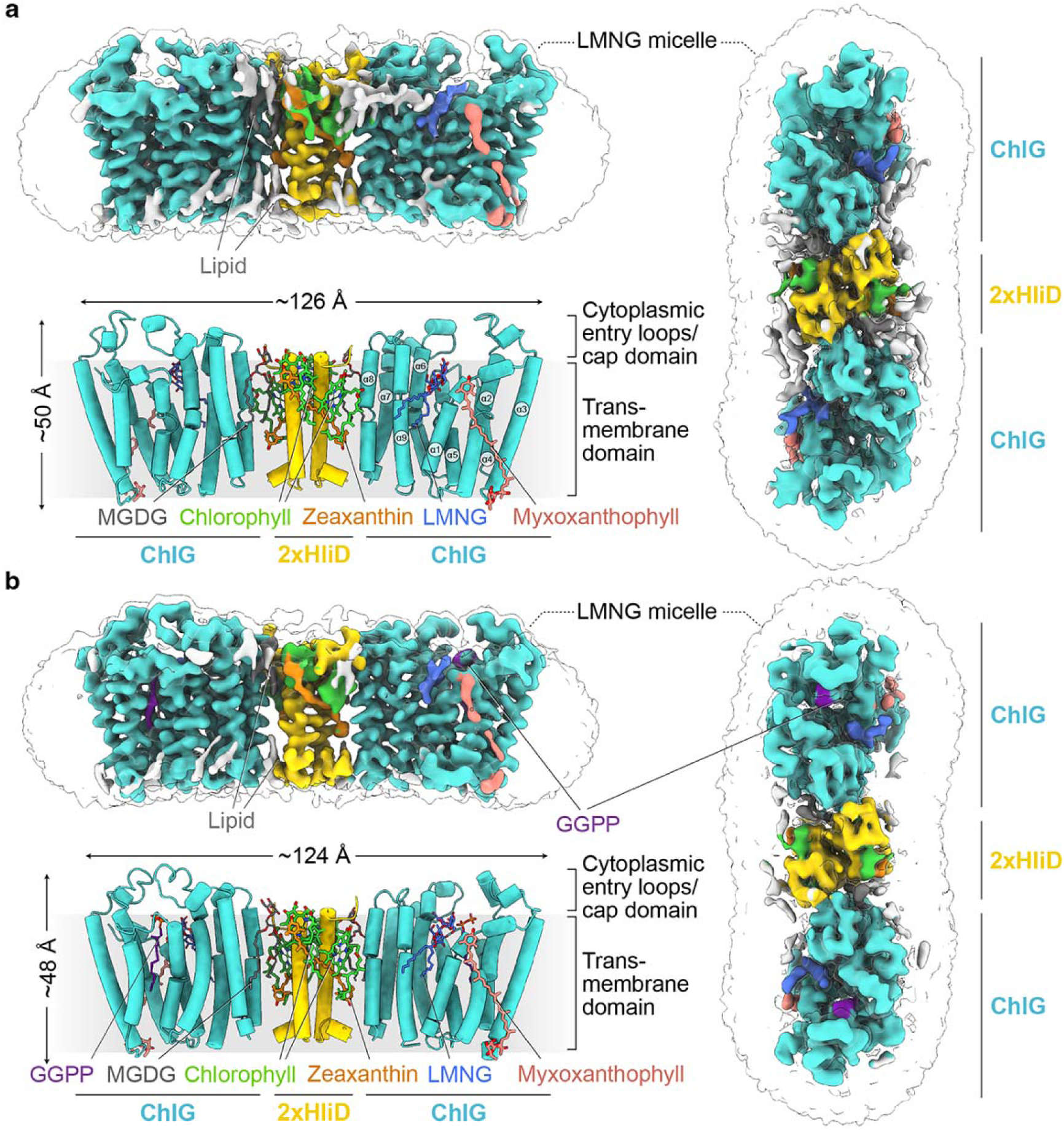
Architecture of the G-D_2_-G complex from *Synechocystis*. **a,** Left, cryo-EM map (top) and structure in cartoon representation (bottom) of the apo G-D_2_-G complex. Right, the cytoplasmic face of the G-D_2_-G complex cryo-EM map. **b,** Left, cryo-EM map (top) and structure in cartoon representation (bottom) of the GGPP-bound G-D_2_-G complex. Right, cryo-EM map of the GGPP-bound G-D_2_-G complex viewed as in (a). Light blue, ChlG; gold, HliD; light green, Chl; orange, Zea; salmon pink, Myx; purple, GGPP; dark grey, monogalactosyldiacylglycerol (MGDG); light grey, lipids. The lauryl maltose neopentyl glycol (LMNG) micelle is shown as a transparent envelope. ChlG α-helices are numbered.

Consistent with its transmembrane nature, the G-D_2_-G complex is nearly fully embedded within the detergent micelle (**Figure 1**), corresponding also to the distribution of its hydrophobic surface (**Figure 2a**). Despite its extended linear shape, the G-D_2_-G complex shows only limited flexibility (**Extended Data Figure 7**). The ChlG monomer contains nine TMHs that create a large active site with two substrate binding domains surrounded by α-helices 1, 2, 4 and 5 in the transmembrane domain. The cavity is gated by cytoplasmic entry loops formed by residues 94-115 and 222-231, which form a cap domain (**Figures 1,2, Extended Data Figure 8**) similar to other prenyltransferases ^10,11,25,26^. The two halves of the complex, each consisting of one ChlG and one HliD, are almost identical to one another, with root-mean-square-deviation (RMSD) values of 1.8 Å (apo) and 2.1 Å (GGPP-bound) (**Extended Data Figure 7d**). In the cryo-EM map of the G-D_2_-G complex incubated with GGPP, a defined extended density located between α-helices α2, α4 and α5 was present in the substrate-binding cavity of each ChlG monomer. This cavity was empty in the apo complex (**Figure 1b, Extended Data Figure 6a**). We attributed this density to a molecule of GGPP, which fitted entirely into this volume (**Figure 2b, Extended Data Figure 6c**).

**Figure 2.**
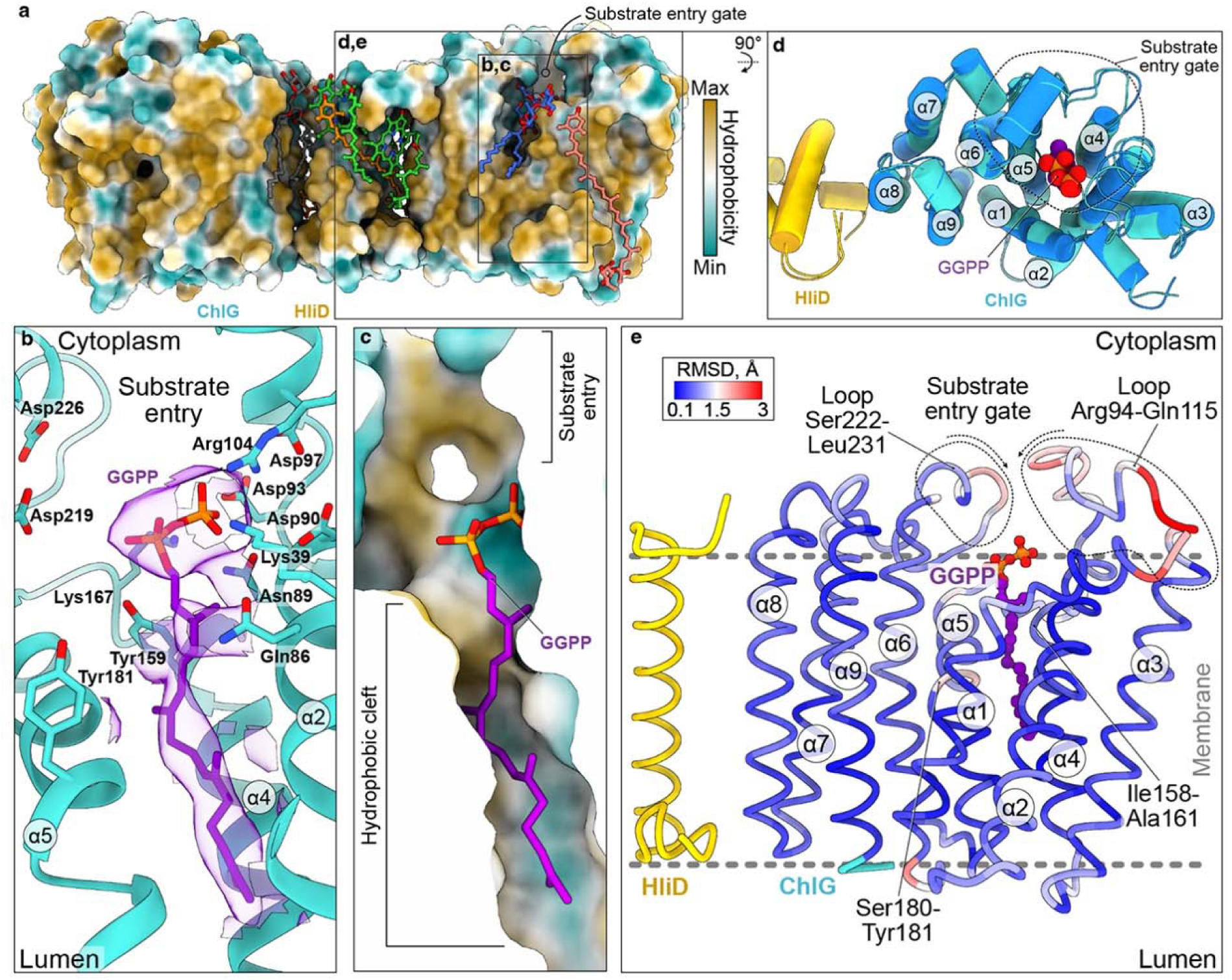
Binding of GGPP to ChlG. **a,** Structure of G-D_2_-G with protein molecules in surface representation colored by hydrophobicity (scale on right) and with molecules of Chl (light green), Zea (orange) and Myx (salmon pink). **b,** Close-up view of the bound molecule of GGPP (purple) with the associated cryo-EM density zoned around it (semi-transparent purple). The amino acid residues of ChlG (light blue) that surround GGPP and are involved in interacting with it (e.g. Asp93, Asn89, Lys39, Arg104) or in catalysis (Tyr159 and Tyr181) are indicated (see also Table 1 and Figure 3). **c,** Same view of bound GGPP as in **(b)**, with the surface of the substrate-binding pocket of ChlG shown and colored by hydrophobicity, as in **(a)**. **d,** Structures of ChlG (light blue) and HliD (yellow) from the GGPP-bound G-D_2_-G complex in cartoon representation (colored) superimposed with the structures of ChlG (dark blue) and HliD (orange) from the apo G-D_2_-G complex (semi-transparent), viewed from the cytoplasmic side. ChlG α-helices are numbered. The bound molecule of GGPP is shown in sphere representation. **e,** Structures of ChlG and HliD (yellow) from the GGPP-bound G-D_2_-G complex in ribbon representation with ChlG colored by the root mean square deviation (RMSD) from the apo G-D_2_-G structure. ChlG α-helices are numbered. The bound molecule of GGPP (purple) is shown in ball and stick representation. The approximate position of the membrane is shown by the dashed grey lines.

**Table 1.** Testing the roles of residues predicted to be important for substrate binding or catalysis in *Synechocystis* ChlG.

| ChlG residue | Proposed role in substrate binding/catalysis | Activity of Ala mutant relative to Ile44Phe ChlG | Photoheterotrophic growth |
| --- | --- | --- | --- |
| Lys39 | GGPP interaction | N.D. | - |
| Gln86 | GGPP interaction | N.D. | - |
| Gln89 | GGPP interaction (via Mg<sup>2+</sup> coordination) | N.D. | - |
| Asp90 | GGPP interaction | N.D. | - |
| Asp93 | GGPP interaction (via Mg<sup>2+</sup> coordination) | N.D. | - |
| Asp97 | GGPP interaction | N.D. | - |
| Arg104 | GGPP interaction | N.D. | - |
| Tyr159 | Catalysis (carbocation stabilisation) | N.D. | - |
| Lys167 | GGPP interaction | N.D. | - |
| Tyr181 | Catalysis (carbocation stabilisation) | 1±0% | - |
| Asn218 | Chlide interaction (C17 carboxyl) | N.D. | - |
| Asp219 | GGPP interaction (via Mg<sup>2+</sup> coordination) | N.D. | - |
| Lys221 | Chlide interaction (C13 ester) | N.D. | - |
| Asp226 | GGPP interaction (via Mg<sup>2+</sup> coordination) | N.D. | - |
| Gln301 | Chlide interaction (C13 ester/ C17 carboxyl) | 101±60% | + |
| Gln305 | Chlide coordination (central Mg) | 13±8% | + |

Opposite to the substrate cavity, ChlG α-helix 8 (α8, residues 268-287) provides extensive contacts to the HliD dimer (see below and **Figures 1,4**), which bridges to the second ChlG monomer on the other side of the complex. In the centre of the complex, each HliD polypeptide interacts with two molecules of Chl and one molecule of Zea (**Figures 1,4**), consistent with high-performance liquid chromatography (HPLC) analysis of pigments in the G-D_2_-G preparation (**Extended Data Figure 2**). Interestingly, we found additional densities at the periphery of the complex, located at the ChlG helices α1-α3, which we attributed to Myx, and which is present in the complex in a similar concentration as Zea (**Extended Data Figure 2**). Additionally, in each half of the complex we found two densities at the cytoplasmic and lumenal sides between ChlG and HliD, which we assigned as lipid molecules (**Figures 1,4b**). The size and shape of the lipid densities on the cytoplasmic side closely resemble the most abundant thylakoid lipid monogalactosyldiacylglycerol (MGDG), which we modelled there (**Extended Data Figure 6b,c**), and seem to provide supplementary hydrophobic interfaces to additionally “glue” ChlG, HliD, Zea and Chl molecules together (discussed further below). In addition, at the lateral entrance into the ChlG active site we found densities that we assigned as lauryl maltose neopentyl glycol (LMNG) molecules (**Figure 1, Extended Data Figure 6b,c**).

Apart from G-D_2_-G, we identified G-D_2_ protein complexes in our cryo-EM samples; despite their relatively low predicted molecular mass of ∼48 kDa, we refined their maps to 6.8 Å (apo) and 6.1 Å (GGPP-bound) (**Extended Data Figure 9**). Although the rather low resolution of the maps did not allow for model building, we were able to assign corresponding densities to a ChlG monomer and a HliD dimer, demonstrating an overall similar architecture to G-D_2_ within the larger G-D_2_-G complex.

### GGPP binding to ChlG

The comparison of the apo and GGPP-bound structures did not reveal pronounced differences in the transmembrane domains, but the GGPP-bound complex appears slightly compacted, with each ChlG monomer located closer to the HliD dimer (**Movie S1**). The RMSD values between the ChlG monomers in the GGPP-bound and apo structures are 1.9 Å and 1.8 Å, respectively, indicating that there are minimal structural differences between them (**Figure 2e**). The pyrophosphate moiety of the bound GGPP is coordinated by several amino acid residues, which are conserved among prenyltransferases and mainly located in the entry (cap) loops (**Figure 2, Extended Data Figure 8**). Asp93 and Asp97 of the conserved D_93_xxxD_97_ motif (**Extended Data Figure 8b**), together with Asn89 and Asp90, appear to stabilize the pyrophosphate, presumably via interactions with a Mg^2+^ ion, as in other prenyltransferases ^10,11,25,26^). Although we could not unambiguously assign densities for Mg^2+^ in our structure, these ions were available in the purification buffer, and the equivalent Asn residue in the *Arabidopsis* ChlG enzyme (Asn155) has previously been shown to be essential for activity in *in vitro* assays ^27^. In addition, the conserved basic Lys39, Arg104 and Lys167 residues form extra electrostatic interactions with the charged pyrophosphate head group of GGPP (**Figure 2b**). Interestingly, Asp226 on the second entry loop (222-231) of the cap domain ^10,11,25,26^ is located too far (about 10 Å) to bind GGPP via coordinated Mg2+, although its conserved counterpart binds the substrate in other prenyltransferases ^10,11,25,26^. The tetraprenyl group of GGPP fits into the cryo-EM density and is anchored deep in one of the substrate-binding cavities of ChlG, surrounded by a number of hydrophobic residues (**Figure 2b,c**).

The cap domain loops show the greatest differences between the apo- and GGPP-bound ChlG structures (**Figure 2e**), suggesting that the flexibility of the cap allows GGPP to enter and bind the enzyme. To elucidate the mechanism of substrate capture, we analyzed the flexibility of the entry loops by MD simulations of ChlG based on our apo and GGPP-bound structures of the complex in the presence of cyanobacterial thylakoid lipids (see Methods). In general, the MD simulations demonstrated rather small fluctuations of the entire ChlG structure, as indicated by slight root mean square fluctuations (RMSF) (**Extended Data Figure 7**). However, the regions within the cap domain (residues 95-106 and residues 220-231) were the most flexible parts of ChlG in both apo and GGPP-bound MD runs (**Extended Data Figure 7a,b**). The average structure of ChlG obtained by MD was also similar to the structure obtained by cryo-EM (**Extended Data Figure 7c**). Interestingly, during the MD simulations, a molecule of MGDG was able to penetrate the ChlG active site through the lateral gate and occupy the position similar to that of LMNG in our cryo-EM structures (**Figure 1**, **Extended Data Figure 7b)**.

To reveal possible conformational variability in the apo and GGPP-bound datasets, we applied the 3D variability analysis (3DVA) tool of CryoSPARC ^28^ (**Extended Data Figure 7e,f**). Using 3DVA we detected only small continuous flexing of the G-D_2_-G complex without any major conformational rearrangements. Again, most of the changes were confined to the vicinity of the substrate-binding pocket, including the cap loops and adjacent helices α2, α3 and α4 (**Extended Data Figure 7e,f**, compare yellow and blue volumes), supporting our MD simulation data.

### Modelling Chlide binding to ChlG

We were unable to obtain a structure of ChlG with its second substrate, Chlide, so to uncover more about its binding mode we performed modelling of *Synechocystis* ChlG with Chlide, GGPP and Mg^2+^ ions using AlphaFold3 (AF3; **Figure 3**). The predicted models agree with the observed placement of the GGPP in the structure obtained by cryo-EM, and the structure at large with an RMSD of 1.69 Å. The models also highlight the predicted location of two Mg^2+^ ions that likely co-ordinate the pyrophosphate group of GGPP and are required for the activation of its electrophilic carbon. The Mg^2+^ ions are coordinated by some of the conserved residues discussed above (Asn89 and Asp93), as well as Asp219 and Asp226 from the second substrate entry gating loop and the following helix α6, which is shifted towards the bound substrates by around 4 Å compared to the cryo-EM structure (**Figure 3a,b**). The AF3 model also positioned Chlide with its esterifiable carboxyl group in proximity to the GGPP activated carbon, where nucleophilic attack from the Chlide carboxyl group is predicted to occur (see Discussion). See **Tables S2** and **S3** for confidence scores (ipTM) of the interaction between the model of ChlG and substrates and for RMSD of Chlide between models. All AF3 calculations can be found in **Extended Dataset S1**.

**Figure 3.**
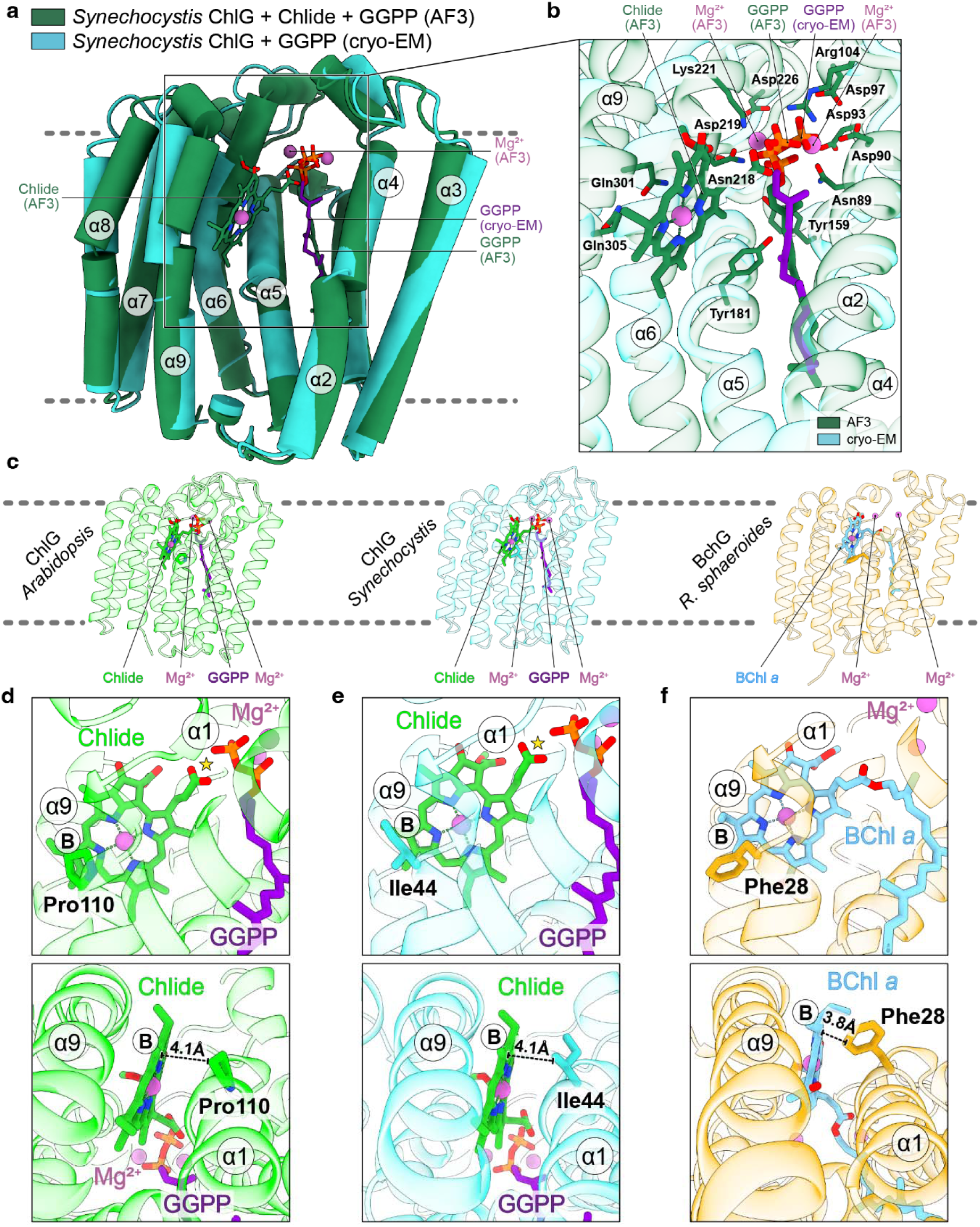
Modelling of Chlide binding to ChlG. **a,** Comparison of the GGPP-bound ChlG structure determined by cryo-EM (light blue) with an AlphaFold3 (AF3) model of ChlG (dark green) with Chlide (green) and GGPP (purple) (both in stick representation) and Mg^2+^ ions (magenta). ChlG α-helices are highlighted and helix α1 is not shown for clarity. **b,** Close-up view of the bound Chlide from **(a)** (prediction, dark green), GGPP (prediction, dark green; cryo-EM data, purple) and Mg^2+^ ions (magenta) in the superimposed models of ChlG obtained by cryo-EM (semi-transparent, light blue) or predicted by AF3 (semi-transparent, dark green). Residues that coordinate Chlide, GGPP, Mg^2+^ ions and/or are important for catalysis are indicated in the AF3 model (also see Table 1). ChlG α-helices are highlighted. **c,** Comparison of AF3 models of plant (*Arabidopsis*, colored light green) and cyanobacterial (*Synechocystis*, colored light blue) ChlG with a purple bacterial BchG (*R. sphaeroides*, colored sand) with bound substrates (or product in the case of BchG). **d-f,** Close-up views of the active sites of the models from (**c**) viewed from two different angles. ChlG and BchG helices α1 and α9 are indicated. The helix α1 residues implicated in substrate specificity are highlighted, and distances from them to Chlide/BChl *a* are shown [Pro110 in plant ChlG (**d**), Ile44 in cyanobacterial ChlG (**e**), and Phe28 in BchG (**f**)]. Ring B of the Chlide/BChl *a* macrocycle is marked. The yellow star indicates the site of GGPP attachment to Chlide (in **d** and **e**).

The modelling highlighted several previously unidentified Chlide-interacting residues that target the C13 and C17 groups of the macrocycle. Asn218 is predicted to form a hydrogen bond with the C17 carboxyl group, ensuring its proximity to the activated carbon of GGPP and its correct orientation for nucleophilic attack. Lys221 provides a hydrogen bond to an oxygen of the C13^2^ ester group, providing additional binding energy and orienting the macrocycle. Gln301 interacts with both the C17 carboxyl group and the C13^2^ ester group. Finally, Gln305, conserved in ChlG enzymes, appears to be proximal to the central magnesium of the Chlide ring and could act as an axial ligand, ensuring further stabilization of substrate binding (**Figure 3a,b**). The model further indicates several previously identified residues conserved in prenyltransferases. In addition to those discussed above two aromatic residues, Tyr159 and Tyr181, located close to the activated carbon on GGPP, are thought to stabilize the δ^+^ charge on the terminal GGPP carbon.

To verify the predicted importance of some of the residues identified above, we screened a series of point mutations using the Ile44Phe variant of *Synechocystis* ChlG that can synthesize BChl in a Δ*bchG* mutant of the purple phototrophic bacterium *R. sphaeroides* (see ^29^ and **Extended Data Figure 10** for more information). Our structure shows that Ile44 is found just after the bend in helix α1 at the lateral intramembrane gate, which we expect provides a connection between the ChlG catalytic site and the lipid environment for substrate entry to the substrate-binding cavity of ChlG and for the product release, as suggested for homologous enzymes ^30^ (**Figure 3c,e**). The amino acid at the corresponding position in an AF3 model of BchG is Phe28 (**Figure 3c,f**), thus this residue must at least partly confer substrate specificity on these enzymes, with the replacement of isoleucine by phenylalanine allowing ChlG to accommodate and esterify BChlide (see *Discussion* section for further explanation).

We tested 16 residues predicted to be required for substrate binding and/or catalysis based on our structural and modelling data by substituting them with alanine and quantifying BChl production compared to the ChlG Ile44Phe ‘parent’ when expressed in a *R. sphaeroides* Δ*bchG* mutant as a proxy for activity (**Table 1** and **Extended Data Figure 10**). When normalized to cell dry weight, all but two of the ChlG variants synthesized ≤ 1% the amount of BChl produced by the Ile44Phe control, only one of which produced detectable quantities of BChl (1%), confirming their importance for activity. This was further supported by the inability of these 14 mutated enzymes to support photoheterotrophic growth of the Δ*bchG* mutant. The exceptions were the Gln301Ala and Gln305Ala variants of ChlG, which retained 101±60% and 13±8% the activity of the Ile44Phe variant based on the BChl level and were both able to support photoheterotrophic growth. We predicted these two glutamines are involved in the binding of the Chlide substrate (**Figure 3b**), one via central Mg coordination (Gln305) and one via hydrogen bond interaction with C13 keto ester oxygen (Gln301). Our results suggest that they are not strictly required for ChlG activity and may represent some redundancy in the binding of the Chlide substrate.

### HliD-pigment subcomplex and its interaction with ChlG

In cyanobacteria, ChlG binds tightly to HliD and the stability of ChlG *in vivo* is compromised in the absence of HliD ^13,21^. Our structure revealed that the X-shaped HliD dimer links two monomers of ChlG (**Figure 4**). Apart from connections through Zea molecules (see below), ChlG monomers interact with the central HliD-pigment subcomplex via C- and N-terminal regions of helix α8 (**Figure 4b,c**). At the lumenal N-terminus of the helix, the interaction is established via hydrophobic residues and a possible hydrogen bond between ChlG Gln268 and HliD Trp53. At the cytoplasmic C-terminus of ChlG helix α8, electrostatic interactions are established through ChlG Asp286 and HliD Arg30.

**Figure 4.**
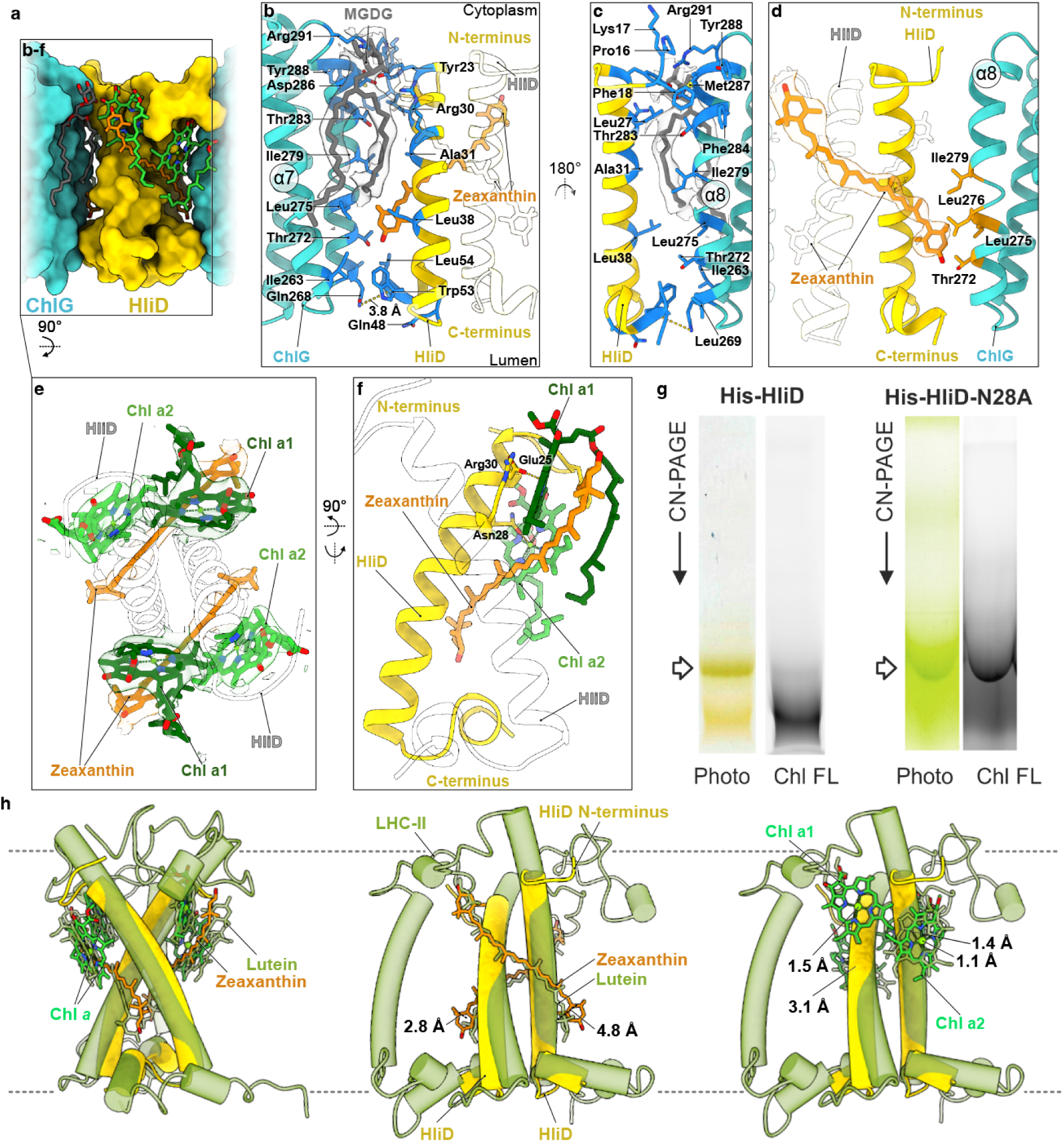
Characterization of HliD-pigment complex and its association with ChlG. **a,** The apo G-D_2_-G complex structure with the models of bound Chls (green), Zea (orange) and Myx molecules (salmon pink). **b,** Close-up view of the interface between ChlG and HliD. Amino acid residues of ChlG and HliD involved in interactions are colored blue; the α-helices of ChlG are indicated and numbered. Semi-transparent cryo-EM densities (grey) of bound monogalactosyldiacylglycerol (MGDG) lipids and molecules of Zea (orange) are shown. Only one of the HliD proteins is colored (yellow). **c,** Close-up view of the HliD-ChlG interface as in **(b)** observed from the opposite side of the complex. Amino acid residues of ChlG involved in interactions with Zea are colored orange. Semi-transparent cryo-EM density is zoned around the molecule of Zea, α-helices of ChlG are indicated. Only one of the HliD proteins is colored (yellow). **d,** The HliD dimer from the apo G-D_2_-G complex observed from the cytoplasmic side. The models of bound pigments with respective semi-transparent cryo-EM densities zoned around are shown (Chl a1/2, dark/light green, Zea, orange). **e,** The HliD dimer from the apo G-D_2_-G complex viewed from the ChlG interface side. Ligand molecules are shown only at one side of the HliD dimer (Chl a1/2, dark/light green, Zea, orange) and only one of the HliD proteins is colored (yellow). HliD residues coordinating bound molecules of Chl are shown. **f,** Comparison of the HliD dimer from the apo G-D_2_-G complex (yellow) with the structure of LHC-II (light green, PDB:1RWT) observed from different sides. Molecules of bound pigments are indicated. Distances between HliD Zea and LHC-II lutein molecules (middle panel), as well as between Chl molecules (right panel) are indicated. The approximate position of the membrane is indicated by grey dashed lines. **g**, HliD and HliD-N28A pulldowns, loaded for a similar concentration of Chl (∼ 1 μg), were separated by CN-PAGE and Chl fluorescence measured using an LAS 4000 imager (Fuji). Arrows mark the dimeric HliD and HliD-Asn28Ala; see the **Extended Data** Fig. 11c for the separation in the second dimension.

In the cryo-EM maps of both the apo and GGPP-bound complexes, we detected clear densities for the chlorin rings of four Chls and two carotenoids on the cytoplasmic side of the HliD dimer (**Figures 1, 4d-f, Extended Data Figure 6b,c**). Due to the flexible nature of the Chl phytyl tails and the limited resolution of the map in this region, we could not model phytyls unambiguously, so we fitted them to the most suitable densities. The two Zea molecules are in a cross-brace arrangement, each located inclined to the membrane plane, with their polyene chains bound to the hydrophobic middle part of the crossed HliD dimer supercoil (**Figure 4a-f**). Their lumenal β-rings are buried between HliD and ChlG and establish contacts with ChlG α8 hydrophobic residues and Thr272 (**Figure 4d**)_;_ MD simulation has shown the presence of hydrogen bond between Zea and Thr272 in 78 % of MD time. The Zea stromal β-rings are located more peripherally and are stabilized by the interactions with the tetrapyrrole ring of Chl a1 (see below) from one side and its phytyl chain from the other (**Figure 4e,f**). The Chl molecules bound to HliD are coordinated through a Chl-binding motif (ExxNxR) conserved in the whole LHC superfamily ^16^. Two Chl molecules near to the HliD N-terminus (referred to as Chl a1) correspond to the 602 and 610 Chls in eukaryotic LHCs (nomenclature from ^31^) and are coordinated by the Glu25 residue from this motif, with Arg30 from the motif of the opposite HliD monomer appearing to interact via pi-cation interactions with the delocalised pi-orbital of the macrocycle (**Figure 4f**). Chls placed closer to the lumenal side of the complex, referred to as Chl a2 and corresponding to the 603 and 612 Chls in LHCs, are coordinated by Asn28 (**Figure 4f**).

Although it is accepted that Hlips are ancestors of the eukaryotic LHC superfamily ^16^, the overall structural similarity of pigments bound to dimeric HliD and LHCs is remarkable. While LHCs bind more Chls and carotenoids than Hlips, there are few differences in the arrangement of the central four Chls (a1/602, a2/603, a1/610, a2/612) and two xanthophylls (L1, L2 sites) in the canonical plant LHC-II (PDB: 1RWT, ^31^) when compared with dimeric HliD (**Figure 4h, left panel**). The individual positions of Chls are shifted by only several Å in respect to the supercoiled HliD dimer, and, surprisingly, while Chl a1 retains the orientation of the corresponding Chl 602/610, Chl a2 is flipped and rotated compared to the corresponding Chl 603/612 in LHC-II (**Figure 4h, right panel, Extended Data Fig. 14**). Alternative orientations of Chl a2 that more closely resemble the positioning of this Chl molecule in the LHC-II complex did not result in a better fit into the cryo-EM map **(Extended Data Fig. 14b,c)**; therefore, Chl a2 was fitted in a flipped and rotated orientation **(Extended Data Fig. 14a)**. Zea molecules in the ChlG-HliD complex are also only slightly shifted, however their β-rings are rotated such that they are almost perpendicular with respect to their counterparts in LHC-II (**Figure 4h, middle panel**).

Chl 612 is typically the lowest-energy Chl in LHCs and is in close contact with the L2 xanthophyll ^32,33^. This Chl is therefore proposed to be the pigment that directs excitation energy to the quenching carotenoid molecule ^32,33^. To probe the role of the corresponding Chl a2 molecules in HliD, we produced wild-type HliD and a mutated Asn28Ala variant as His-tagged proteins in *Synechocystis*. Because HliD can form also heterodimers with another Hlip HliC, we further deleted the *hliC* gene to isolate pure homodimeric HliD complexes. The HliD and HliD-N28A preparations were separated on a CN gel loaded at a comparable level of Chl and the intensity of Chl fluorescence of wild-type HliD and the Asn28Ala variant was then analyzed in the gel. While the wild-type protein showed almost no fluorescence as expected for the quenched Chls in Hlips ^20^, the Asn28Ala variant was brightly fluorescent (**Figure 4g**).

We eluted both dimeric complexes from the native gel and measured their absorbance spectra, which revealed a lower carotenoid content for the Asn28Ala variant than for the wild-type protein (**Extended Data Figure 11a)**. The shifted Chl to β-Car ratio was confirmed by HPLC (**Extended Data Figure 11b**); the content of β-Car (0.22) per Chl in the Asn28Ala complex was much lower than the ∼0.5 β-Car per Chl ratio for wild-type HliD, which was also reported previously ^15,20^. Although free HliD binds β-Car, not Zea, we expect the binding site to be the same (^22^; see following discussion). Further separation of the isolated complex in the second dimension by SDS-PAGE showed a much higher amount of Asn28Ala HliD (**Extended Data Figure 11c**) than wild-type HliD, although the loaded Chl concentration was comparable. This indicates that a large fraction of the mutated protein either binds no pigments or pigments were lost during purification. The presence of Chl a2 ligated by HliD Asn28 is thus critical for the binding of remaining pigments (Chl a1 and carotenoid) in a stable configuration.

## Discussion

### The mechanism of Chlide prenylation and substrate specificity of ChlG and BchG enzymes

ChlG belongs to the UbiA superfamily of intramembrane prenyltransferases that exhibit conserved folding, but members of the superfamily recognize chemically distinct prenyl acceptor substrates and different isoprenyl chains ^10,11,25,26^. Indeed, ChlG shares a high sequence similarity with other prenyltransferases and adopts a conserved nine-TMH architecture (**Extended Data Figure 8**). Here, we determined the cryo-EM structure of ChlG with and without the GGPP substrate, revealing the isoprenyl substrate binding mode (**Figures 1, 2**). The overall structural rearrangement of ChlG after binding of GGPP is not large, similar to the homologous archaeal UbiA protein ^10^. However, our cryo-EM data, supported by MD simulations and structural modelling predictions (**Figure 2,3, Extended Data Figure 7**), suggest that the observed flexible movement of the cytoplasmic substrate entry loops of the cap domain facilitate GGPP gaining access to the ChlG binding cavity from the surrounding membrane through the lateral gate between helices α1 and α9. This is consistent with MD simulations, in which a molecule of the MGDG lipid was able to penetrate from the membrane and occupy the enzyme active site (**Extended Data Figure 7b**), representing a potential mechanism for ChlG substrate entry. After GGPP enters ChlG, the entry loops adopt a more closed conformation (**Figure 2d,e**), interacting with the pyrophosphate group of GGPP to stabilize the bound substrate within the cavity. Indeed, mutation of the residues in this region abolished the activity of ChlG (**Table 1**). Following esterification of the Chlide macrocycle, GG-Chl *a* product likely exits the active site through the same lateral gate used for substrate entry. It remains unclear where and how the geranylgeranyl tail of Chl *a*_GG_ is reduced by ChlP ^2^, as well as how the synthesized Chl molecule is transferred to photosynthetic machinery proteins ^34^.

We were unable to obtain a Chlide-bound structure of ChlG, which is not surprising given the expected instability of the ChlG-Chlide complex, as Chlide binding requires preloading of ChlG with prenyl pyrophosphate ^35^. Therefore, to investigate the binding of Chlide to ChlG, we used structure predictions by AF3 ^36^ in combination with site-directed mutagenesis. The predicted topology of Chlide bound to ChlG favors prenylation with its carboxyl group oriented towards the bond between the pyrophosphate and isoprenyl moieties of GGPP (**Figure 3b**), with the carboxyl oxygen only 4.3 Å from the terminal GGPP carbon. This is consistent with previous data obtained with chemically modified Chlide derivatives. While plant ChlG tolerates bulky substituents on ring B, which is facing the open cavity lateral gate of ChlG in our model, the enzyme did not accept substrates with bulky substituents on rings C and E ^37^, which are deeply buried inside the substrate binding cavity (**Figure 3b**).

Based on our structures, modelling and activity data, the catalytic mechanism of ChlG is likely similar to other members of the UbiA superfamily ^10^, although due to the strong nucleophilic nature of the carboxyl oxygen compared to the standard prenyltransferases, it may proceed via an S_N_2 rather than S_N_1 reaction. Additionally, the prerequisite of GGPP binding before Chlide binding and the inherent instability of a carbocation required for the S_N_1 reaction that would likely form before Chlide binding suggests the concerted S_N_2 reaction. Either way, this results in the formation of an ester bond between Chlide and the GG tail (**Figure 5a**). Although we could not clearly assign Mg^2+^ ions in our cryo-EM density, we expect them to be essential for the enzymatic activity based on our structural modelling (**Figure 3**), mutational analysis (**Table 1**) and the homology of ChlG to other prenyltransferases ^10,11,25,26^. As both GGPP and phytyl-PP and can serve as substrate for ChlG ^6^, we modelled the structures of *Synechocystis* ChlG, *Arabidopsis* ChlG and *R. sphaeroides* BchG with both Chlide and either GGPP or phytyl-PP (**Extended Data Figure 13**). The models show nearly identical substrate positioning in both cases, suggesting that the reaction would proceed in the same way.

**Figure 5.**
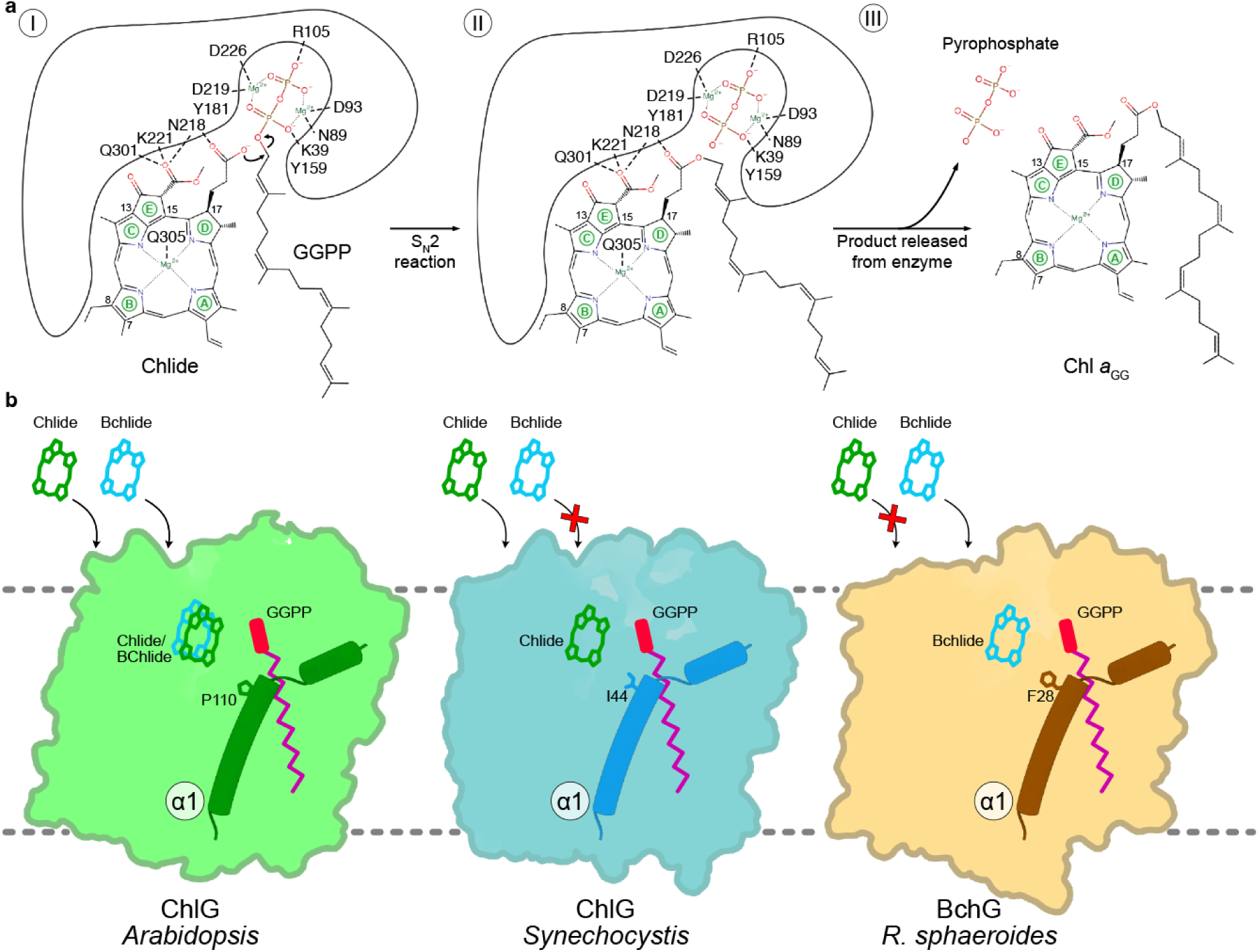
Mechanism of catalysis and substrate selectivity of ChlG and BchG enzymes. **a**, Proposed mechanism of the ChlG-catalyzed attachment of the GG tail to Chlide, following an S_N_2 reaction resulting in the formation and release of Chl *a*_GG_ and pyrophosphate. Macrocycle rings and selected carbon atoms of Chlide/Chl *a* macrocycle are marked. **b**, Schematic comparison of plant (light green) and cyanobacterial (light blue) ChlG with BchG from purple bacteria (sand). The Chlide/BChlide specificity is at least in part provided by the different residues in helix 1 (α1) located close to the substrate entry site, which facilitate the binding of the preferred substrate.

Although Chl synthases possess a certain flexibility for their prenyl donor and acceptor substrates ^37,38^, they do not prenylate protochlorophyllide (Pchlide), an earlier intermediate in Chl biosynthesis ^39^. This selectivity is most likely provided by the presence of Tyr181 in close proximity (about 4.9Å) to the C17-C18 single bond of Chlide (**Figure 3b, Extended Data Figure 12**), which is unreduced and less flexible in PChlide and cannot tolerate such a bulky residue nearby.

The discrimination between Chlide and BChlide, which differ only in the C3 position (vinyl versus acetyl group) and the oxidation state of the C7-C8 bond (single versus double) (**Extended Data Figure 1a**), seems to be at least in part mediated by the residue at the lateral gate of the enzyme (position 28 in *R. sphaeroides* BchG and 44 in *Synechocystis* ChlG, ^29^) which is positioned proximal to the B ring of the (B)Chlide macrocycle (**Figure 3d-f**). The bulky Phe residue makes BchG specific for BChlide, which is relatively more flexible than Chlide due to the tetrahedral rather than trigonal geometry of its C8 carbon. *Synechocystis* ChlG has Ile44 at the equivalent position to Phe28, while the ChlG enzymes from angiosperms, which can also esterify BChlide to some extent ^40^, have a proline residue (Pro110 in *Arabidopsis* ChlG). Thus, having a large aromatic Phe at the lateral gate position in BchG appears to prevent the undesirable esterification of Chlide, an intermediate in BChl biosynthesis pathway that must undergo BChl-specific modifications to rings A and B prior to the addition of the tail (**Extended Data Figure 1)**. The smaller hydrophobic Ile makes cyanobacterial ChlG specific for the less flexible Chlide substrate, and the comparatively smaller and rigid Pro allows the angiosperm enzyme to work on both Chlide and BChlide, although the latter substrate is not relevant *in planta* (**Figure 3d, 5b**). A second level of selectivity may involve the C3-acetyl group of BChlide, which is a vinyl group in Chlide; in BchG Glu157 is positioned proximal to the modified A ring of BChlide, whereas both *Synechocystis* and *Arabidopsis* ChlG have an isoleucine residue at this position (Ile182 and Ile241, respectively, also see **Extended Data Figure 12**). In fact, AF3 models of I44F and WT ChlG with BChlide and Chlide predict that BChlide binds non-productively in WT ChlG but productively in I44F ChlG (**Extended Data Figure 15**). Nevertheless, future structural studies on substrate binding to ChlG and BchlG are necessary to fully clarify the substrate selectivity in these enzymes.

### HliD and the non-substrate pigments attached to ChlG

In early studies, a complicated mixture of carotenoids associated with the isolated ChlG complexes was difficult to interpret ^13,14^. In particular, the presence of both Zea and β-Car was confusing, as free HliD dimers bind β-Cars only (**Extended Data Figure 11b**) ^18^. Our recent work clarified that the isolation of ChlG via an engineered FLAG-tag results in a mixture of several ChlG complexes, namely the G-D_2_-G, the smaller G-D_2_ complex, and ChlG associated with a HliD-HliC heterodimer (G-D/C) ^22^. These complexes bind different carotenoids, which was unexpected. Variants with only one copy of ChlG (G-D_2_, G-D/C) bind Zea and β-Car in a 1:1 ratio whereas the G-D_2_-G contains Zea and only a very low level of β-Car (**Extended Data Figure 2b**) ^22^.

The G-D_2_-G structure supports our previous model that the ‘default’ carotenoid binding to HliD is β-Car unless this protein associates with ChlG. In this case, the same binding site is occupied by Zea because the Thr272 residue of ChlG creates a hydrophilic patch in the vicinity of Zea hydroxyl group. As the G-D_2_-G complexes are destabilised in the *Synechocystis* mutant cells that contain β-Car but lack Zea ^21^, the interaction between helix α8 of ChlG and the Zea seems to play an important structural role in the complex, probably serving as a ‘glue’ at the HliD-ChlG interface. The affinity of β-Car to the HliD-ChlG interface should be therefore significantly weaker than Zea. This explains why the G-D_2_-G assembly, with each HliD copy in contact with ChlG, binds two Zea, while the G-D_2_ complex is purified with one Zea and one β-Car ^22^. While we detected G-D_2_ particles in our cryo-EM datasets, the resolution achieved for the refined maps was not sufficient to identify the bound carotenoids (**Extended Data Figure 9a,b**). However, the presence of G-D_2_, but not G-D, particles in the samples suggests that a hypothetical ChlG complex with only a single copy of HliD is either not stable or the ChlG always binds to a pre-formed Hlip dimer.

Myx is another carotenoid associated with the ChlG complex (**Extended Data Figure 2b**) and cryo-EM data revealed a weak density, close to the lateral gate of the ChlG catalytic side, that matches a molecule of Myx (**Figure 1, Extended Data Figure 6b,c**). This is consistent with the 77K ultrafast spectroscopy measurement indicating that Myx is located at the periphery of ChlG and that there is no energy transfer between this carotenoid and the HliD-associated Chls ^14^. It is worth noting that specific loss of Myx does not affect the interaction of ChlG and HliD ^21^. The role of Myx is not clear but photoprotection of the enzyme seems the most plausible. Although Myx is located too far (∼ 13Å) from the substrate binding site to quench a potential triplet state of Chlide during catalysis, it could quench triplets in Chl products following release from the synthase active site and/or singlet oxygen. While we could not confidently determine the position of the sugar moiety, we placed it facing the lumenal side based on the most convincing fitting into the cryo-EM density (**Extended Data Figure 6b,c**), however, we cannot exclude an alternative orientation of Myx.

### Non-photochemical quenching of HliD chlorophylls

The energy absorbed by the Chl molecules in the G-D_2_-G complex is very rapidly transferred to the Zea molecules and dissipated as heat in about 20 ps via the internal conversion of the carotenoid ^14^. This mechanism involves a direct energy transfer from the Q_y_ of the excited (singlet) Chl to the optically forbidden S1 state of the carotenoid ^14^. Basically, the same mechanism of quenching seems to be present in the whole LHC superfamily including the LHC antennas ^41^ ^42^.

Based on resonance Raman spectroscopy measurements, it has been proposed that specific pigment−protein or pigment−pigment interactions lower the carotenoid S1 energy, for instance by twisting carotenoid molecules ^43^. Unexpectedly, there is no obvious mechanism lowering the energy of Zea in the G-D_2_-G structure; based on our cryo-EM map, both Zea molecules appear to be in the all-*trans* configuration and are not substantially bent or twisted. However, due to the relatively small size of the G-D_2_-G complex and its slight fluctuations (**Extended Data Figure 7e**), we could not achieve sufficient resolution to precisely assign the configuration of Zea molecules and their distances from the Chls. However, in comparison with luteins in LHC-II (**Figure 4h**), Zea molecules are rotated 90° along their polyene chain with their β-rings positioned at a different angle towards Chl a1. In addition, the chlorin ring of Chl a2 in HliD is flipped and also rotated counterclockwise by around 90° compared to Chl 603/612 of LHC-II (**Figure 4h, Extended Data Figures 14,15**). Moreover, the distance between Chl a2 and Zea seems to be closer (∼ 4.5-5 Å) than the corresponding Chl 603/612-lutein distance (∼ 5.5-6 Å) in the crystal structure of LHC-II, which already exhibits a reduced fluorescence lifetime (∼1 ns) relative to detergent-solubilized LHC-II complexes (3-4 ns) ^31^.

As discussed recently ^33^, even a slight difference in the distance between 612-type Chl and carotenoid could have significant impact on the Chl Q_y_ to carotenoid S1 energy transfer. It is therefore possible that the relative distance together with other factors (e.g. carotenoid rotation) allow fast quenching in Hlips. Removing Chl a2 abolished quenching in HliD but also compromised the stability of the remaining pigments. Consistent with our previous work on the LHC-like protein LIL3 ^41^, it is very likely that all three pigments (Chl a1/2 and a carotenoid) must bind to at least one Hlip helix to form a stable and quenched pigment assembly.

## Methods

### Synechocystis strains and cultivation

All strains used in this study were constructed in a *Synechocystis* GT-P background ^44^. The *f.chlG*/Δ*chlG*/Δ*hliC*/Δ*ycf39* strain was described in ^22^ and the *psbAII*:*his-hliD*/Δ*hliC*/Δ*hliD* strain was generated by deleting the *hliC* gene in the *psbAII*:*his-hliD*/Δ*hliD* strain ^45^ as described in ^20^. To obtain the *psbAII*:*his-hliD-N28A*/Δ*hliC*/Δ*hliD* strain the *psbAII*:*his*-*hliD*-Km^R^ plasmid ^45^ was modified by QuikChange II XL site-directed mutagenesis kit (Agilent Technologies) to replace the HliD-Asn28 residue with alanine using primers HliD-N28A-f: GGCAGCCCGGCCAGCGAGCTTTTCCGCGT and HliD-N28A-r: ACGCGGAAAAGCTCGCTGGCCGGGCTGCC). The resulting *psbAII*:*his*-*hliD-N28A*-Km^R^ plasmid was transformed to the Δ*hliD* mutant ^13^ and the mutated *psbAII*:*hliD-N28A* locus was fully segregated by plating on an increasing concentration of kanamycin, followed by the deletion of *hliC* as described above.

For the purification of the G-D_2_-G complex, an 8 L of the *Synechocystis f.chlG*/Δ*chlG*Δ*hliC*/Δ*ycf39* strain was grown to an optical density at 750 nm (OD_750_) of 1.0-1.5 in a homemade bioreactors bubbled with sterile air at 28 °C under 40 µmol photons m^-2^ s^-1^. For the isolation of His-HliD and His-HliD-N28A, *psbAII*:*his-hliD*/Δ*hliC*/Δ*hliD* and *psbAII*:*his-hliD-N28A*/Δ*hliC*/Δ*hliD* cells were cultivated in 1 L cylinders with 700 mL BG11, at 28 °C and under 40 µmol photons m^-2^ s^-1^.

### Isolation of protein complexes and 2D protein electrophoresis

The isolation and solubilization of thylakoid membranes by a mixture of *n*-dodecyl-β-D-maltoside and glycol-diosgenin was described in ^22^. The G-D_2_-G complex was purified on an anti-FLAG column as described in ^24^, however a mixture of 0.02% (w/v) *n*-dodecyl-β-D-maltoside and 0.02% (w/v) glycol-diosgenin in thylakoid buffer (25 mM MES, pH 6.5, 10 mM MgCl_2_, 10 mM CaCl_2_, 25% glycerol) was used instead of 0.04% (w/v) *n*-dodecyl-β-D-maltoside for all steps of the purification. The purification of His-tagged HliD and HliD-N28A was performed essentially as described in ^20^. The isolated protein complexes were separated by CN-PAGE with 1% (w/v) A8-35 amphipol according to ^46^. Individual components of the protein complex were resolved by incubating the gel strip from the first dimension in 2% (w/v) SDS and 1% (w/v) dithiothreitol for 30 min at room temperature, and proteins were separated in the second dimension by SDS-PAGE in a denaturing 16-20% (w/v) polyacrylamide gel containing 7 M urea ^47^. Proteins were stained with Coomassie Brilliant Blue.

### Absorption spectra and analysis of pigments

Absorption spectra of purified protein complexes were measured as described in ^20^. To analyse the pigments associated with protein complexes, specific bands from CN gels containing the respective protein complexes were excised, followed by pigment extraction and detection by high-performance liquid chromatography (HPLC) as described in ^20^.

### Cryo-EM sample preparation and data acquisition

3 µL of the isolated G-D_2_-G complex at a protein concentration of approximately 2 mg mL^-1^ was applied onto double-glow-discharged C-Flat grids (R1.2/1.3 3Cu-50) (EMS) and immediately plunge-frozen in liquid ethane using a Vitrobot Mark IV (Thermo Fisher Scientific) with the environmental chamber set at 100% humidity and 4 °C. For the GGPP-bound sample, the protein complex was purified in the presence of 10 mM Mg^2+^ and incubated with 150 µM of GGPP (Sigma-Aldrich) for 30-40 min at room temperature prior to plunge-freezing. Data acquisition (22,594 movies for the apo sample and 18,660 movies for the GGPP-bound sample) was performed automatically using a Glacios cryogenic transmission electron microscope (Thermo Fisher Scientific) operating at 200 kV with a Selectris imaging energy filter and a Falcon 4i direct electron detector (both Thermo Fisher Scientific). Data collection was performed in Electron Event Representation mode at a nominal magnification of 165,000x (0.68 Å per pixel) in the defocus range of −0.8 to −1.8 μm, with an exposure time of approximately 3.9 s for the apo samples, and 4.3 s for the GGPP-bound sample, resulting in a total electron dose of 50 e^-^ Å^−2^.

### Cryo-EM data processing

Data analysis workflows for the apo and GGPP-bound samples were similar with minor variations (for details on the cryo-EM processing workflow, see **Extended Data Figures 3-5, 9**). Cryo-EM data quality assessment and initial steps of processing were performed in cryoSPARC Live, with further processing carried out in cryoSPARC v4 and v5 ^48^. Motion correction, contrast transfer function (CTF) estimation, primary rounds of particle picking and 2D classification (using Fourier-cropped box) were performed in cryoSPARC Live. Extensive particle picking was continued in cryoSPARC v4 using template and neural network-based Topaz picking pipelines ^49^, followed by rounds of thorough 2D classification of particles. Selected high-quality particles were subjected to duplicate removal and were combined from multiple collected datasets (**Extended Data Figures 4, 5**).

For the apo complex sample, the next processing steps were as follows. 1.266 million particles selected after 2D classifications were extracted using the full box size of 512 pixels and subjected to a round of ab-initio 3D reconstruction with six classes followed by heterogeneous refinement (using the cropped box size of 256 pixels). Two classes, one representing the G-D_2_-G complex [∼365,000 particles, 4.5 Å resolution at a Gold Standard Fourier Shell Correlation (GSFSC) value of 0.143] and the other representing the G-D_2_ complex (∼188,000 particles, 7.3 Å resolution at GSFSC of 0.143), were further processed separately. The G-D_2_-G class was used for three successive rounds of heterogeneous refinement with two “trash” classes to further filter out junk particles, followed by non-uniform (NU)-refinement ^50^ of the best class. The reconstruction obtained after the third round of these refinements (∼354,000 particles, 3.41 Å resolution at GSFSC of 0.143, referred to as NU1) was further subjected to local refinement, NU-refinement and another round of local refinement using a mask covering the entire volume. This map achieved resolution of 3.39 Å (GSFSC of 0.143). In parallel, the NU1 refinement volume was used in another round of heterogeneous refinement with five “trash” classes and using preselected ∼954,000 particles from initial picking to improve the resolution. The resulting good class was used in successive ab-initio 3D reconstruction and heterogeneous refinement with five classes followed by heterogeneous refinement with four classes. The best class from the four-class heterogeneous refinement was used for further NU-refinement followed by Rebalance Orientations and Subset Particles jobs. The resulting particles were subjected to another round of NU-refinement followed by the Global CTF Refinement, additional NU-refinement and Reference Based Motion Correction (RBMC). The resulting particles were used for another NU-refinement (∼210,000 particles, 3.38 Å resolution at GSFSC of 0.143, referred to as NU2), which was further refined with C2 symmetry (∼210,000 particles, 3.22 Å resolution at GSFSC of 0.143, referred to as NU3), subjected to Symmetry Expansion followed by local refinements of the masked ChlG (∼421,000 particles, 3.14 Å resolution at GSFSC of 0.143) and central HliD-dimer (∼421,000 particles, 3.04 Å resolution at GSFSC of 0.143) parts of the volume (**Extended Data Figure 3b-f, 4**). The G-D_2_ complex (∼188,000 particles, 7.3 Å resolution at GSFSC of 0.143) was further subjected to a round of ab-initio reconstruction/heterogeneous refinement with three classes, NU-refinement, additional ab-initio reconstruction with four classes followed by three rounds of heterogeneous refinement with three classes using two “trash” classes (**Extended Data Figure 9c**). Finally, the best class from the last round of heterogeneous refinement (∼140,000 particles, 7.8 Å at GSFSC of 0.143) was subjected to NU-refinement that produced a G-D_2_ map resolved to 6.8 Å (GSFSC of 0.143) (**Extended Data Figure 9a,c**).

For the GGPP-bound sample, the processing workflow was generally similar to that for the apo sample (**Extended Data Figure 5**). 1.368 million particles combined from all collected datasets were extracted with the full box size of 512 pixels and subjected to a round of ab-initio reconstruction and heterogeneous refinement with six classes. Two classes corresponding to the G-D_2_-G (∼311,000 particles, 5.5 Å at GSFSC of 0.143) and G-D_2_ (∼221,000 particles, 7 Å at GSFSC of 0.143) complexes were further processed separately. For the G-D_2_-G complex, the particles were subjected to two rounds of two-class heterogeneous refinement with a “trash” class followed by NU-refinement. Afterwards, local refinement was performed, followed by another two-class heterogeneous refinement with a “trash” class. Finally, another NU-refinement (referred to as NU5, see **Extended Data Figure 5**) followed by local refinement using a mask covering the entire volume resulted in a map of the G-D_2_-G complex resolved to 3.57 Å (GSFSC of 0.143). In parallel, the volume from the NU refinement after the first round of two-class heterogeneous refinement (referred to as NU4) was used in another round of two-class heterogeneous refinement followed by NU-refinement, three-class heterogeneous refinement and another NU-refinement. The volume from this NU-refinement was further processed by heterogeneous refinement with five “trash” classes and using preselected ∼1.255 million particles from initial picking to improve the resolution. The resulting good class was used in successive ab-initio 3D reconstruction and heterogeneous refinement with five classes followed by heterogeneous refinement with four classes. The best class from the four-class heterogeneous refinement was used for further NU-refinement followed by Rebalance Orientations and Subset Particles jobs. The resulting particles were subjected to another round of NU-refinement followed by the Global CTF Refinement, additional NU-refinement and RBMC. The resulting particles were used for another NU-refinement (∼108,000 particles, 3.49 Å resolution at GSFSC of 0.143, referred to as NU6), which was further refined with C2 symmetry (∼108,000 particles, 3.22 Å resolution at GSFSC of 0.143, referred to as NU7), subjected to Symmetry Expansion followed by local refinements of the masked ChlG (∼216,000 particles, 3.27 Å resolution at GSFSC of 0.143) and central HliD-dimer (∼216,000 particles, 3.17 Å resolution at GSFSC of 0.143) parts of the volume (**Extended Data Figure 3h-l, 5**). The G-D_2_ class was further subjected to two-class heterogeneous refinement with a “trash” class followed by NU-refinement, two-class heterogeneous refinement with a “trash” class and another final NU-refinement. The resulting G-D_2_ map was resolved to 6.1 Å (GSFSC of 0.143) (**Extended Data Figure 9b,d**).

Unsupervised B-factor sharpening was applied and local resolution estimation for the final G-D_2_-G maps was generated in cryoSPARC v4. 3D variability analysis ^28^ to investigate the flexibility of the G-D_2_-G complex was performed in cryoSPARC v4 using three different principal components of variability (**Extended Data Figure 7e,f**).

### Model building and refinement

AF3 ^36^ was used to generate initial structural models of the G-D_2_-G complex from *Synechocystis* (Uniprot IDs: P72932 and Q55145). Initial models were manually fitted to the final consensus NU refinement maps of the apo (NU2) and GGPP-bound (NU6) G-D_2_-G complex. Additional maps (Extended Table S1) were used to support model building. Models of Chl *a*, LMNG, MGDG and GGPP molecules were directly added to the models using Coot’s ^51^ function “Get monomer” and models of Zea and Myx were first generated using Grade Web Server tool ^52^ and then added to the models of the complexes in Coot. The models of both the apo and GGPP-bound G-D_2_-G complexes were manually refined in Coot and transferred to Phenix ^53^. The Phenix-implemented ReadySet! program was used to prepare the manually adjusted PDB files of the apo and GGPP-bound G-D_2_-G complexes for further Real Space Refinement procedure in Phenix. Iterative rounds of Phenix Real Space Refinement followed by manual adjustments in Coot were performed to optimise the fit of the models to the cryo-EM maps of the complexes. Validation of the models was done using MolProbity ^54^ in Phenix (see Table S1 for model refinement and validation statistics). Cryo-EM maps and models were visualized in UCSF ChimeraX ^55^ and structural figures were prepared in Affinity Designer 2.

### Molecular dynamics simulations

CHARMM-GUI ^56^ was used for preparation complexes for MD, including ligand parameterization with CGenFF version 3.0 and membrane-embedding. Coordinates models of D_2_-G (two runs, monomers 1, 2 of ChlG) and G-D_2_-G (monomers 3,4 of ChlG) complexes were obtained from the apo G-D_2_-G cryo-EM structure. The membrane lipid content was as follows: SQDG (12%), MGDG (50%), DGDG (20%), POPG (15%), corresponding to the lipid composition of *Synechocystis* cells ^58^. MD simulations were performed for 1 μs for 4 replicates in GROMACS 2022 ^59^ with CHARMM36 force field, V-rescale thermostat and C-rescale barostat were applied. The analysis of MD was done with GROMACS 2022, for visualization xmgrace and visual molecular dynamics (VMD) were used ^60^.

### Structure prediction with AlphaFold3

The AF3 ^36^ model was cloned from GitHub (https://github.com/google-deepmind/alphafold3), which includes instructions for downloading the weights and protein databases. Prediction of the *A. thaliana* ChlG structure was carried out using the sequence from Uniprot (Q38833) and the chloroplast transit peptide deleted (1-57) with Chlide *a*, GGPP and two Mg^2+^ ions. Prediction of the *Synechocystis* sp. PCC 6803 ChlG structure was carried out using the sequence from Uniprot (Q55145) with Chlide *a*, GGPP and two Mg^2+^ ions. Prediction of the *R. sphaeroides* BchG structure was carried out using the sequence from Uniprot (Q9Z5D6) with BChl *a* and two Mg^2+^ ions. The calculations were performed on an 80 Gb NVIDIA A100 GPU.

### Assaying ChlG activity in R. sphaeroides

In contrast to the situation in *Synechocystis*, where *chlG* is essential ^13^, preventing segregation of strains that lack ChlG activity ^61^, which is time consuming and promotes suppressor/reversion mutations, the *bchG* gene (like BChl biosynthesis in general) is dispensable for chemoheterotrophic growth in *R. sphaeroides* ^9^. Expression of the *Synechocystis chlG* gene in a *R. sphaeroides* Δ*bchG* background did not initially result in BChl biosynthesis (**Extended Data Figure 10a)**, but incubation under photoheterotrophic conditions selected for an Ile44Phe suppressor mutation of ChlG that partially restored BChl biosynthesis and growth (**Extended Data Figure 10b,c**), as reported previously ^29^. Due to the difficulty in producing ChlG in sufficient yields for high-throughput *in vitro* assays ^27^, and the aforementioned essentiality of the enzyme in cyanobacteria, we used the Ile44Phe variant in the *R. sphaeroides* Δ*bchG* mutant to test additional ChlG point mutants, using the amount of BChl produced compared to the Ile44Phe ‘parent’ strain and whether each variant strain can grow photoheterotrophically to illustrate the effect on ChlG activity (**Table 1**).

The WT and Δ*bchG* strains of *R. sphaeroides* were grown in M22+ media supplemented with 0.1% (w/v) Casamino acids ^62^. Solid media was set with 1.5% (w/v) bactoagar (Oxoid) and kanamycin (30 µg ml^-1^) was added to plates/cultures where necessary to select for and/or maintain plasmids. Liquid cultures were grown under either semi-aerobic conditions in darkness (chemoheterotrophic growth) or anaerobically with illumination (photoheterotrophic growth), as outlined previously ^63^. The Δ*bchG* strain was generated by allelic exchange using the pK18mob*sacB* plasmid, following the detailed protocol described in Sutherland *et al*. ^64^, such that all but the first 9 and last 6 bp of the gene were deleted and replaced by an XbaI site, which was confirmed by sequencing of the *bchG* locus PCR amplified from the mutant strain (Eurofins Genomics). The *R. sphaeroides bchG* and *Synechocystis chlG* genes were PCR amplified using Q5^®^ Hot Start High-Fidelity DNA Polymerase (New England Biolabs), digested using Thermo Scientific FastDigest restriction enzymes, and ligated (using T4 DNA ligase from New England Biolabs) into the BglII and NotI sites of the pBBRBB-Ppuf843-1200 plasmid ^65^. Point mutants of the gene encoding the Ile44Phe variant of ChlG were generated using the Agilent QuikChange II Site-Directed Mutagenesis Kit according to the manufacturer’s instructions. All plasmids were verified by Sanger sequencing (Eurofins Genomics) prior to introduction to the Δ*bchG* mutant by conjugation ^64^.

To quantify the amount of BChl *a* produced, strains were grown chemoheterotrophically in 80 mL cultures contained within 125-mL Erlenmeyer flasks, harvested by centrifugation (4,200 × g for 30 min at 4°C), and all the supernatant was removed. The pellets were resuspended in 2 mL of 20 mM Tris pH 8 and the wet cell weight was recorded. 100-200 µl of resuspended cells (equating to 10-30 mg) were pelleted (15,000 × g for 5 min at 4°C), the supernatant was removed, and the pigments were extracted from the pellets in 0.5-1 mL of methanol in darkness. Following centrifugation (15,000 × g for 5 min at 4°C), 50 µl of the clarified pigment extracts were analyzed by high-performance liquid chromatography (HPLC) using the program described in ^61^ with elution of BChl *a* detected by monitoring absorbance at 770 nm (**Extended Data Figure 10d**).

## Acknowledgements

This work was supported by project 22-03092S of the Czech Science Foundation and by the Czech Ministry of Education, project PHOTOMACHINES, CZ.02.01.01/00/22_008/0004624. DS acknowledges the support by the German Research Foundation (Emmy Noether grant 537976353, SFB1557) and the Department of Biology’s early-career fellowship at Osnabrück University. We thank Prof. Arne Moeller for access to the cryo-EM platform at the CellNanOs center at Osnabrück University (DFG grant INST190/196-1 FUGG), as well as for his continued support, insightful discussions and useful comments on the paper. KOP and FSM-B were supported by University of Sheffield Faculty of Science PhD studentships. CNH acknowledges Synergy Award 854126 from the European Research Council. AH acknowledges the support of a Royal Society University Research Fellowship (URF\R\241006).

## Data availability

Atomic coordinates have been deposited in the PDB: chlorophyll synthase in complex with the LHC-like protein HliD, apo state (PDB ID code 9SAS), chlorophyll synthase in complex with the LHC-like protein HliD, GGPP-bound state (PDB ID code 9SAU). The associated cryo-EM density maps, have been deposited in the Electron Microscopy Data Bank (EMDB): chlorophyll synthase in complex with the LHC-like protein HliD, apo state, consensus map NU2 (EMD ID code EMD-54696), chlorophyll synthase in complex with the LHC-like protein HliD, GGPP-bound state, consensus map NU6 (EMD ID code EMD-54697), chlorophyll synthase in complex with the LHC-like protein HliD, apo state, consensus map (EMD ID code EMD-54698), chlorophyll synthase in complex with the LHC-like protein HliD, GGPP-bound state, consensus map (EMD ID code EMD-54699), chlorophyll synthase in complex with the LHC-like protein HliD, ChlGx1-HliDx2 complex, apo sample (EMD ID code EMD-54700), chlorophyll synthase in complex with the LHC-like protein HliD, ChlGx1-HliDx2 complex, GGPP sample (EMD ID code EMD-54701), chlorophyll synthase in complex with the LHC-like protein HliD, apo state, full-map local refinement (EMD ID code EMDB-57787), chlorophyll synthase in complex with the LHC-like protein HliD, apo state, consensus map NU3 (C2 symmetry) (EMD ID code EMDB-57781), chlorophyll synthase in complex with the LHC-like protein HliD, apo state, HliD local refinement map (EMD ID code EMDB-57782), chlorophyll synthase in complex with the LHC-like protein HliD, apo state, ChlG local refinement map (EMD ID code EMDB-57783), chlorophyll synthase in complex with the LHC-like protein HliD, GGPP-bound state, full-map local refinement (EMD ID code EMDB-57788), chlorophyll synthase in complex with the LHC-like protein HliD, GGPP-bound state, consensus map NU7 (C2 symmetry) (EMD ID code EMDB-57784), chlorophyll synthase in complex with the LHC-like protein HliD, GGPP-bound state, HliD local refinement map (EMD ID code EMDB-57785), chlorophyll synthase in complex with the LHC-like protein HliD, GGPP-bound state, ChlG local refinement map (EMD ID code EMDB-57786).

## Author contributions

DS, AH and RS designed the study. AW, AP, JP and RS purified and analyzed the ChlG-HliD complexes. DS prepared the cryo-EM grids, collected cryo-EM data, and built and refined the models. DS, FSM-B, CNH, AH and RS analyzed the structure and interpreted the structural data. FSM-B performed structure predictions and NK carried out the MD simulations. KOP, MSP and AH generated and assayed variants of ChlG. The supervision of the project and coordination of writing the manuscript were done by DS, CNH, AH and RS. All authors contributed to writing and proofreading of the manuscript.

