## Extended Fig. S1-S15; Extended Table S1-S3 for "Structure of chlorophyll synthase in complex with the LHC-like protein HliD"

[3] Center of Cellular Nanoanalytic Osnabrück (CellNanOs); Osnabrück University, Osnabrueck 49076, Germany

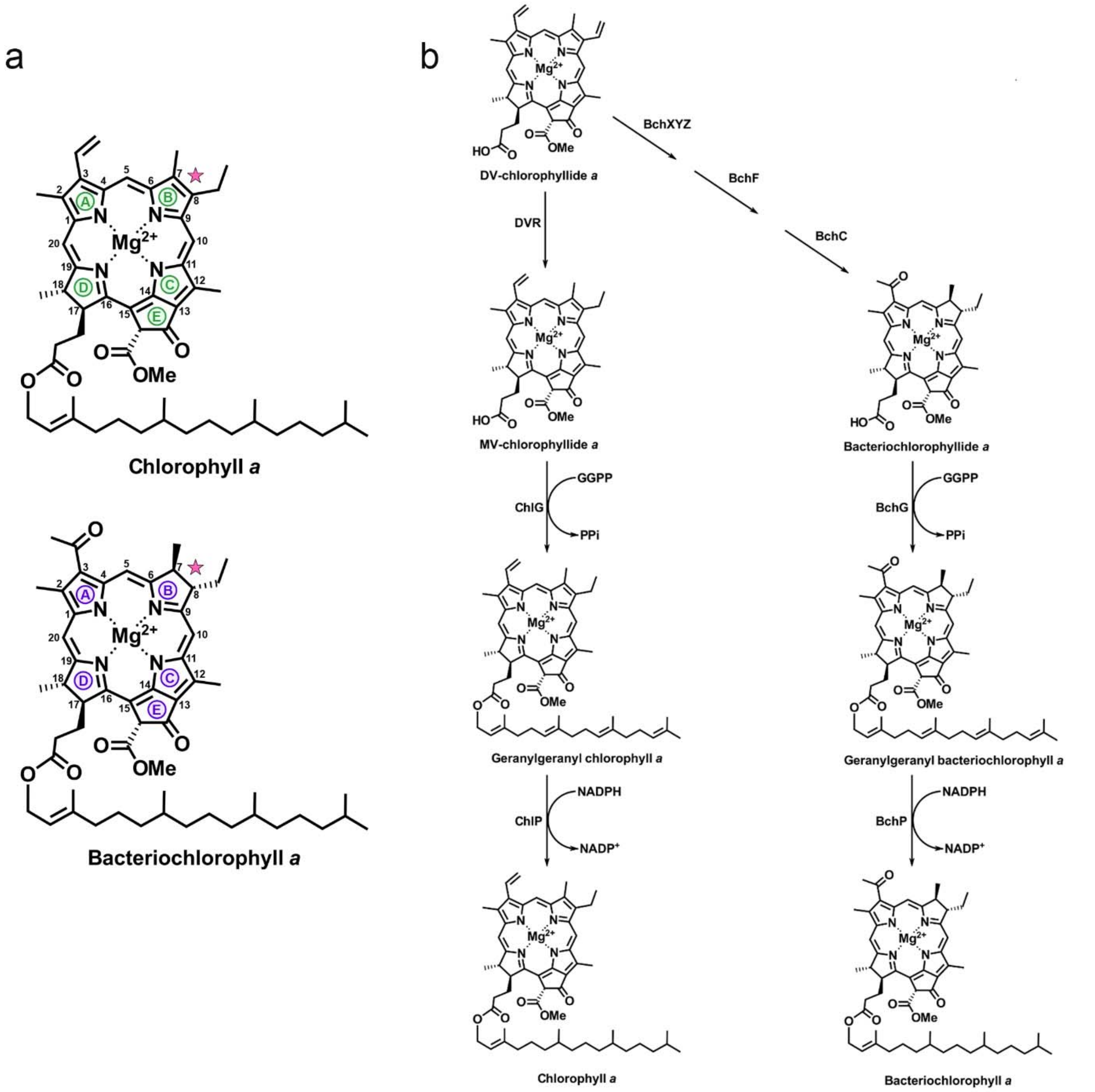


**Extended Data Figure 1. The latter stages of chlorophyll *a* and bacteriochlorophyll *a* biosynthesis.**

**a**, Chemical structures of chlorophyll *a* and bacteriochlorophyll *a*. Carbon atoms and macrocycle rings are indicated. Pink stars highlight the structural differences in the oxidation state of the C7-C8 bond in the ring B between the chlorophyll *a* and bacteriochlorophyll *a* macrocycles. **b**, The biosynthesis of chlorophyll *a* and bacteriochlorophyll *a* follows the same pathway to the point of divinyl (DV)-chlorophyllide *a* (the steps before this intermediate are omitted from this figure). In chlorophyll *a* biosynthesis, the activity of divinyl reductase (DVR) generates monovinyl (MV)-chlorophyllide *a*, the substrate of ChlG, whereas in bacteriochlorophyll *a* biosynthesis chlorophyllide *a* oxidoreductase (COR or BchXYZ) acts on DV-chlorophyllide *a*, with subsequent reactions catalysed by BchF and BchC generating bacteriochlorophyllide *a*, the substrate of BchG.

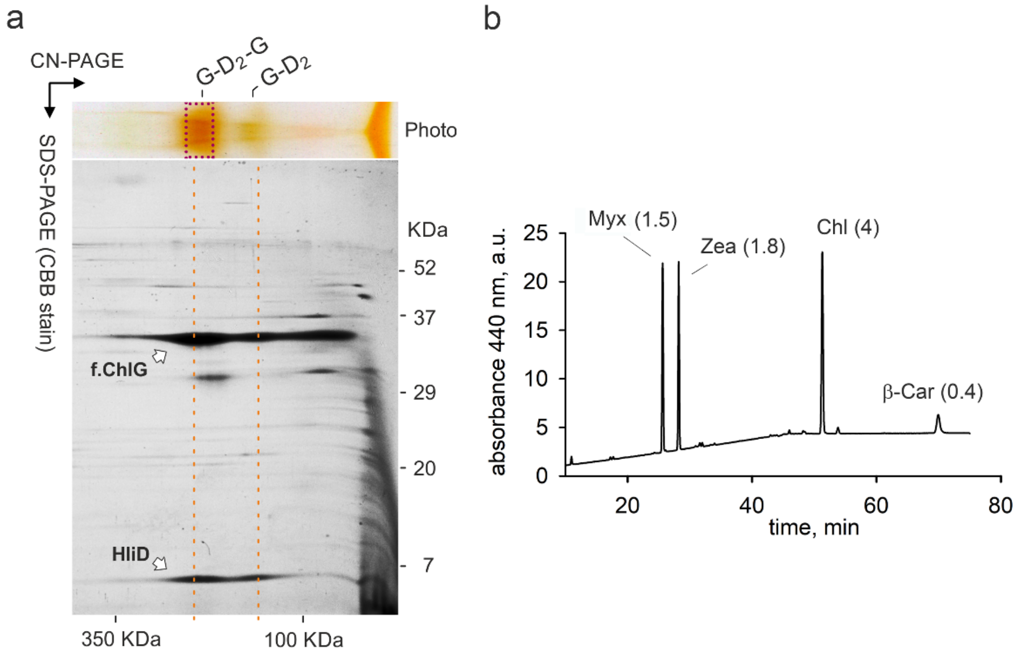


**Extended Data Figure 2. Biochemical characterization of the ChlG-HliD complexes used for cryo-EM**.

**a**, The ChlG-HliD$_2$-ChlG (G-D$_2$-G) complex was purified from the *Synechocystis* FLAG-*chlG/ΔhliC/Δycf39* strain [1] and then separated by clear-native electrophoresis (CN-PAGE). A single strip from CN-PAGE was placed at the top of SDS-PAGE gel and proteins were separated in the second dimension and the gel was stained with Coomassie Blue (CBB stain). The obtained spots were designated as f.ChlG or HliD according to [1]. The preparation also contains a lesser amount of the smaller ChlG-HliD$_2$ (G-D$_2$) complex. **b**, The part of the gel indicated by dashed box in (a) was excised, the protein eluted, and the extracted pigments analysed by HPLC; the relative amounts of myxoxanthophyll (Myx), zeaxanthin (Zea) and β-carotene (β-Car) were calculated per 4 Chl molecules (values in parentheses).

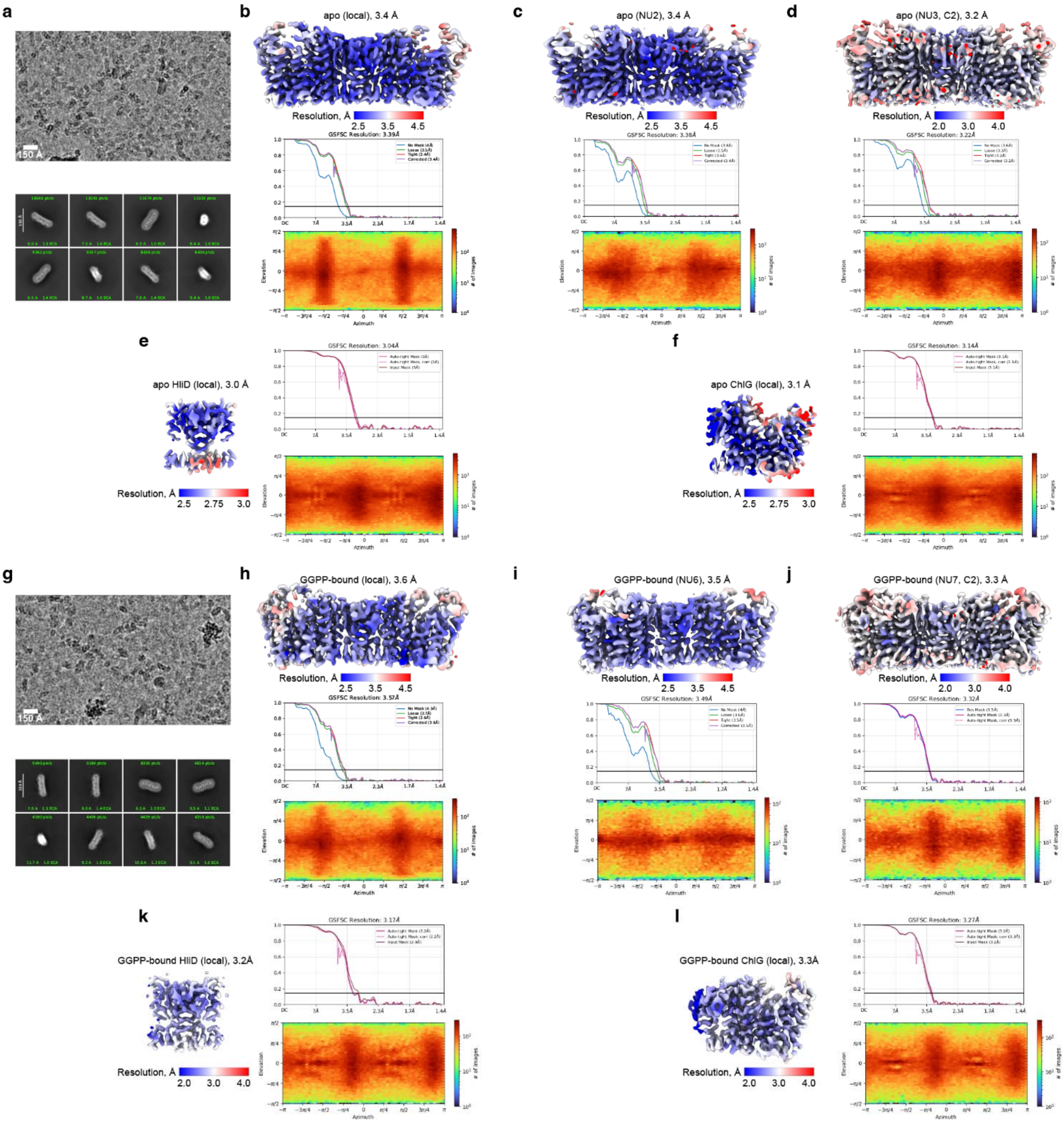


**Extended Data Figure 3. Single particle cryo-EM data analysis and validation for the apo (a-f) and GGPP-bound (g-l) G-$D_2$-G complexes.**

**a,g,** Representative cryo-EM micrograph (top) and 2D class averages (bottom). **b-f** and **h-l,** Local resolution estimation map generated in cryoSPARC (top panels, maps are colored by local resolution); cryoSPARC-generated Gold standard Fourier shell correlation (GSFSC) curves (middle panels); cryoSPARC-generated angular distribution heatmap plots (bottom panels). See methods and Extended Data Fig. 4 and 5 for details.

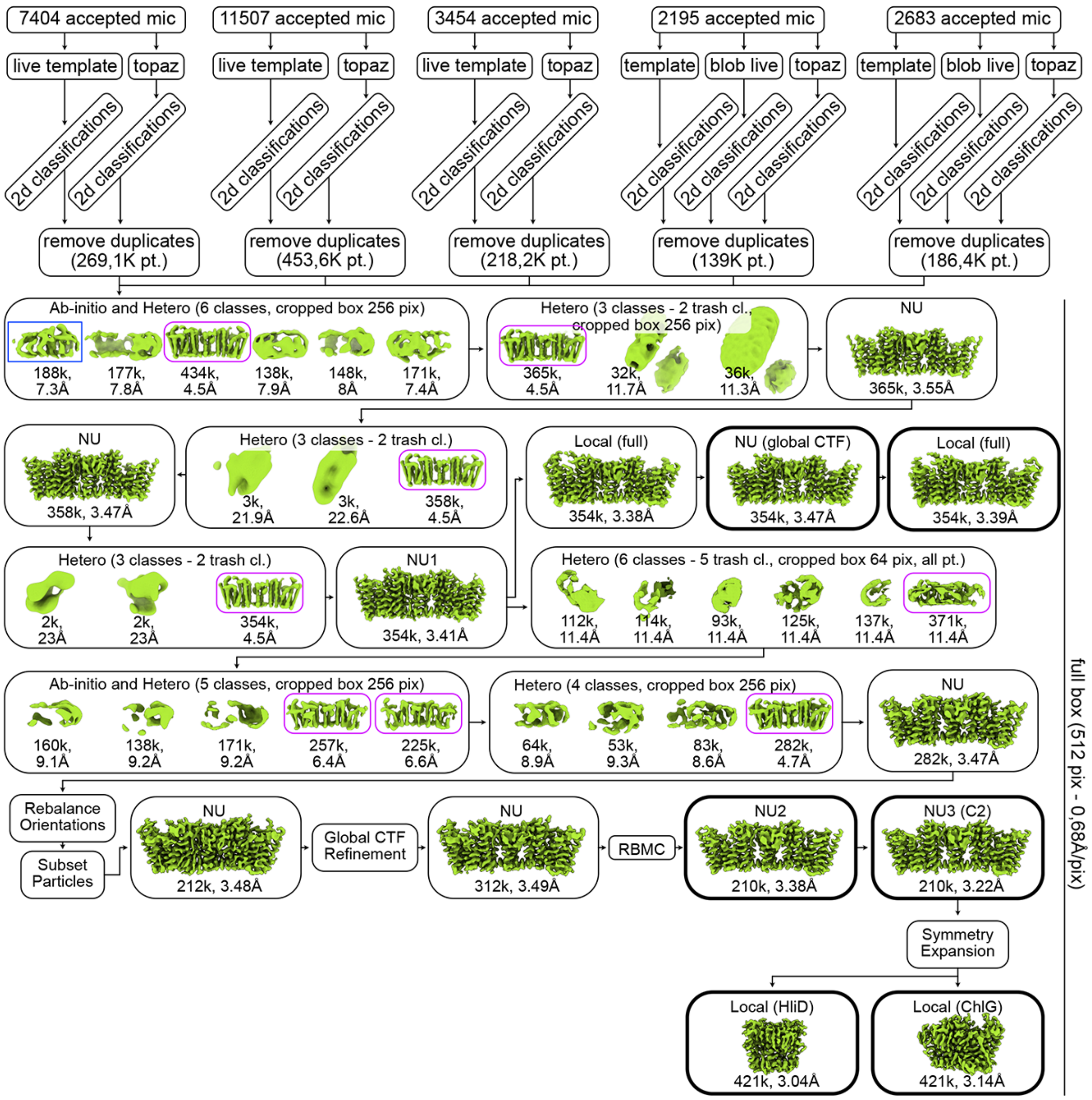


**Extended Data Figure 4. Cryo-EM data processing pipeline scheme for the apo G-D$_2$-G complex.**

To remove duplicates, ab-initio reconstruction, and refinement jobs, numbers of particles used, and resolutions achieved are indicated. Pt., particles, mic, micrographs, pix, pixel, cl., class. NU, non-uniform refinement, CTF, contrast transfer function, RBMC, reference-based motion correction. Purple frames indicate reconstructions used for the G-D$_2$-G processing; blue frame indicates the reconstruction used for the G-D$_2$ processing. The final refinement jobs are highlighted by boxes with a bold outline.

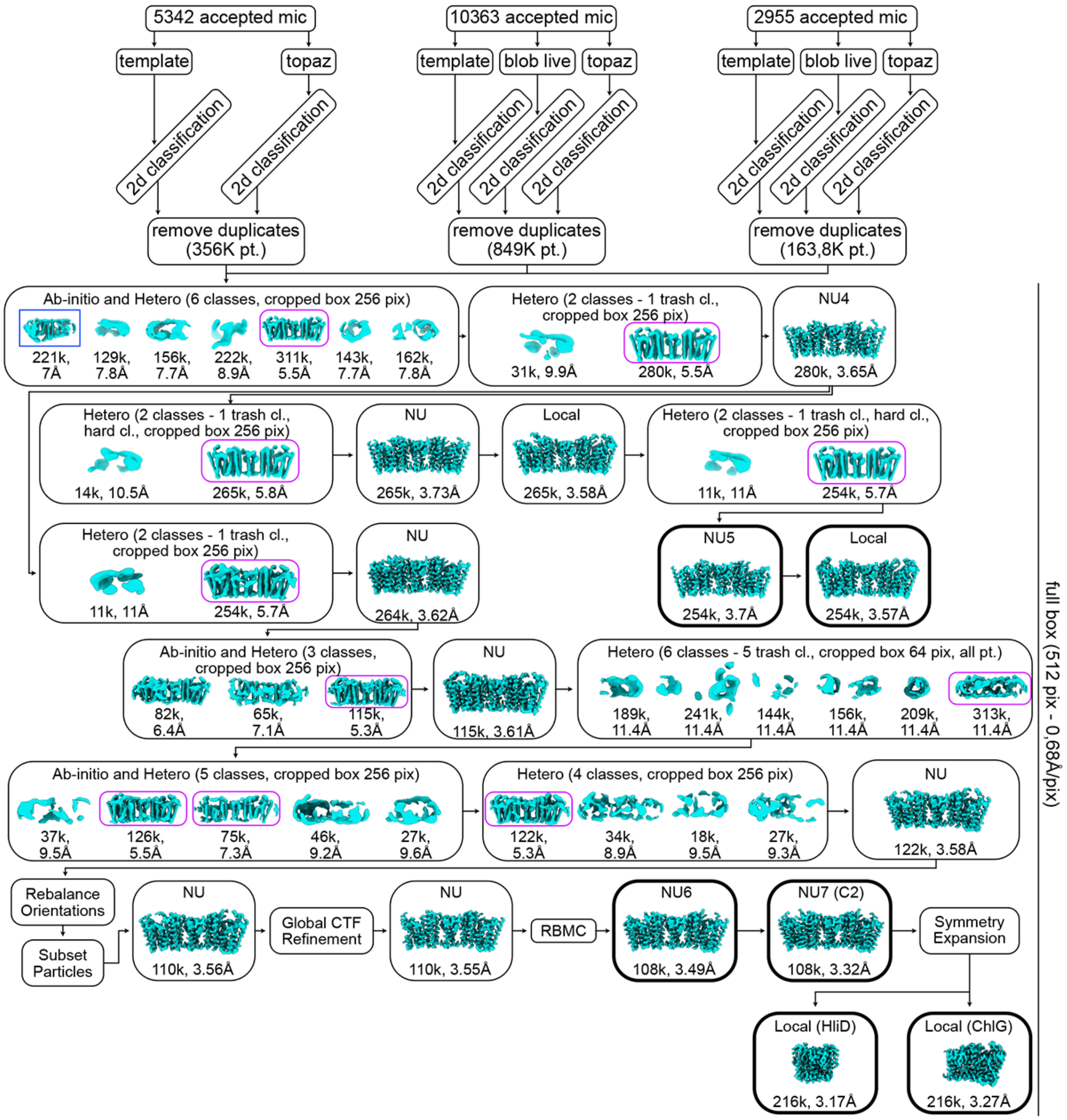


**Extended Data Figure 5. Cryo-EM data processing pipeline scheme for the GGPP-bound G-$D_2$-G complex.**

To remove duplicates, ab-initio reconstruction, and refinement jobs, numbers of particles used, and resolutions achieved are indicated. Pt., particles, mic, micrographs, pix, pixel, cl., class, hard cl., hard classification on. NU, non-uniform refinement, CTF, contrast transfer function, RBMC, reference-based motion correction. Purple frames indicate reconstructions used for the G-$D_2$-G processing; blue frame indicates the reconstruction used for the G-$D_2$ processing. The final refinement jobs are highlighted by boxes with a bold outline.

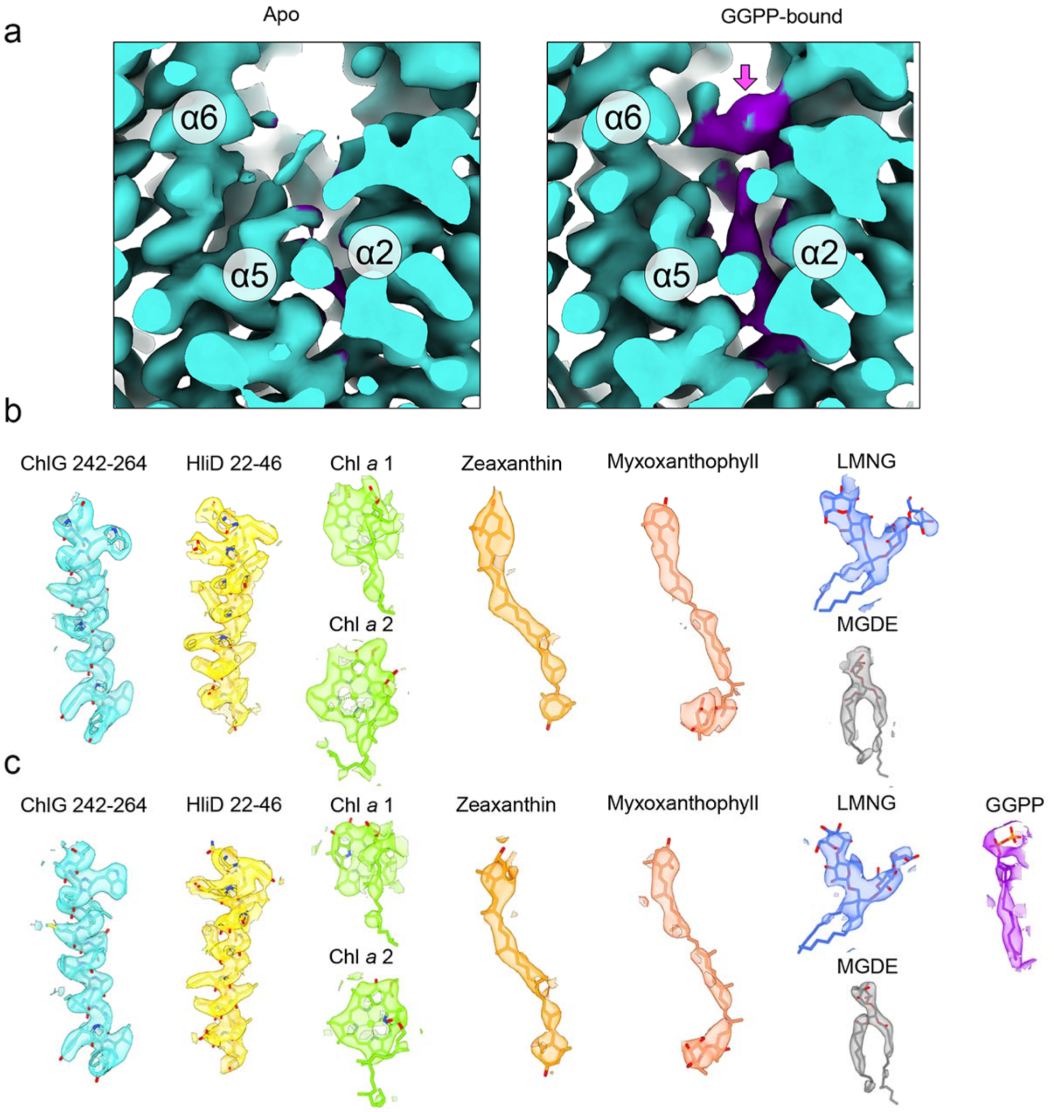


**Extended Data Figure 6. Cryo-EM map quality validation for the G-$D_2$-G complex.**

**a,** Comparison of the cryo-EM map segments representing the substrate-binding pocket of the apo (left) and GGPP-bound (right) G-$D_2$-G complex. α-helices 2, 5, and 6 are indicated. Purple density corresponding to the GGPP molecule is seen in the right panel and is indicated by a pink arrow. **b,c,** Model/map fit of selected protein segments and bound ligands for the final maps of the apo (**b**) and GGPP-bound (**c**) G-$D_2$-G complex.

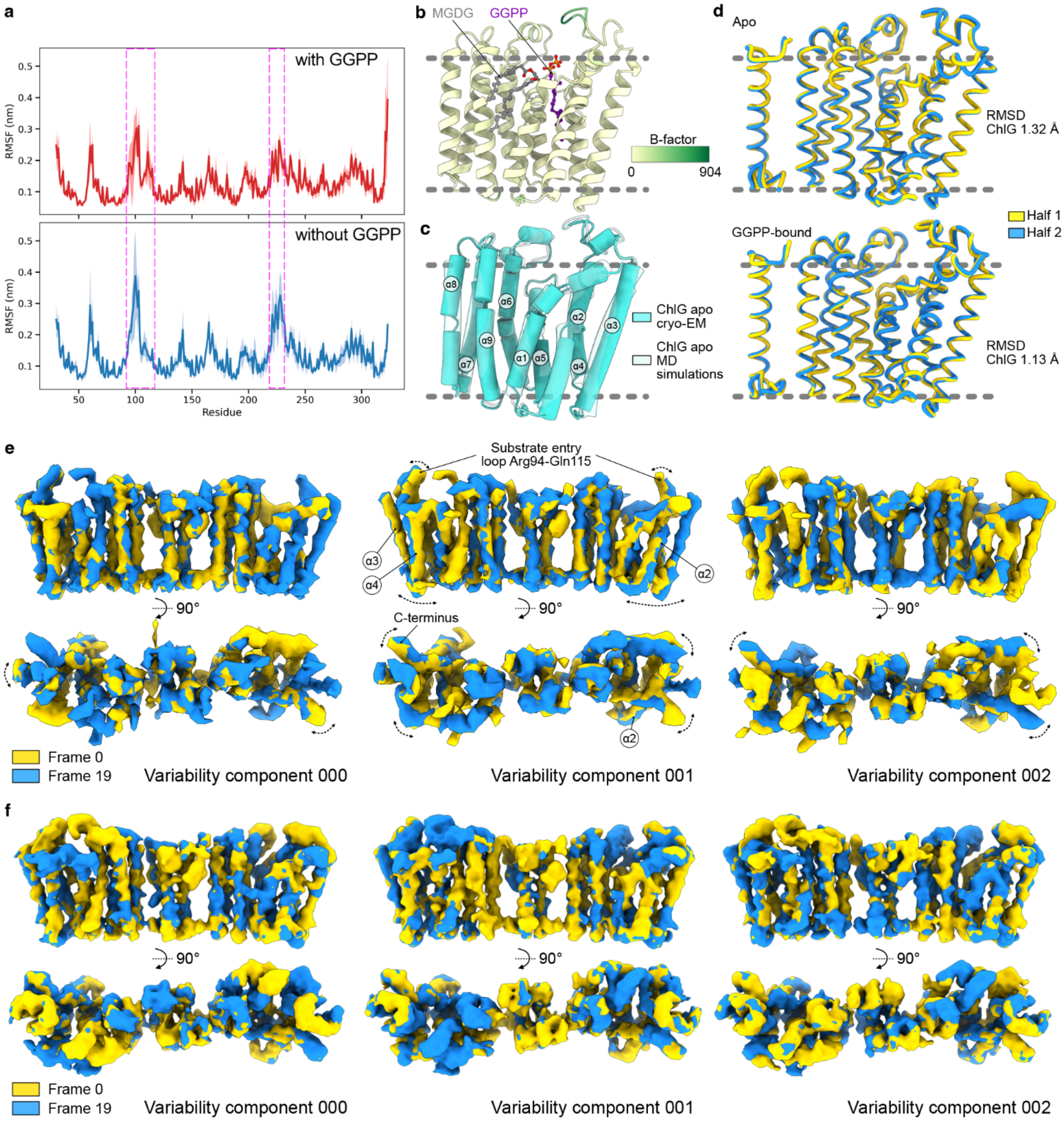


**Extended Data Figure 7. Flexibility of the ChlG and the G-$D_2$-G complex.**

**a**, Root mean square fluctuation (RMSF) of ChlG with or without GGPP (upper panel, red and blue, lower panel respectively) during molecular dynamics (MD) simulations. The calculated RMSF data from four runs was averaged and is shown as solid lines with standard deviation as a shade. The magenta dashed boxes indicate the most flexible regions of ChlG. **b**, Average structure of ChlG without GGPP colored by the B-factor, calculated from MD simulation run 1. A molecule of monogalactosyl diacylglycerol (MGDG, grey) penetrating the substrate-binding cavity of ChlG, as well as a molecule of GGPP taken from our GGPP-bound cryo-EM structure of ChlG are shown. **c**, Superposition of the apo ChlG structure determined by cryo-EM (cartoon representation, solid light blue color) with the structure from the MD simulations without GGPP (semi-transparent, light blue color). ChlG α-helices are numbered. **d**,

Superposition of the opposing halves of the G-$D_2$-G complex in ribbon representation in the apo (top) and GGPP-bound (bottom) state solved by cryo-EM. Root mean square deviation (RMSD) is calculated for C-alpha atoms of ChlG. **e**, Density maps generated by 3D variability analysis (3DVA) for the apo sample at negative (yellow, frame 1) and positive (blue, frame 19) positions along each of the three used variability components. Most variable α-helices of ChlG are indicated. **f**, Density maps generated by 3DVA for the GGPP-bound sample. Coloring is as in panel **d**.

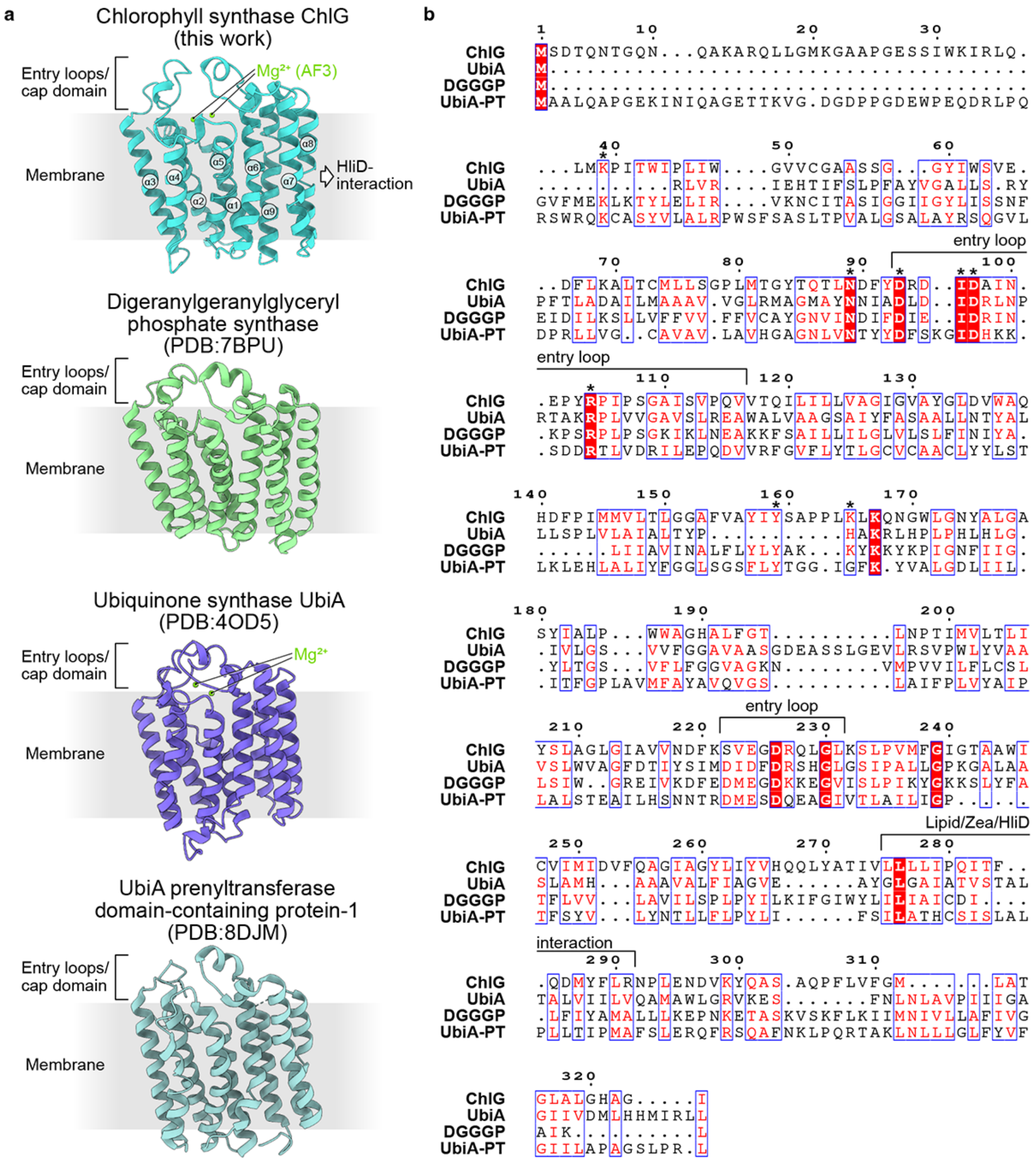


**Extended Data Figure 8. Comparison of *Synechocystis* ChlG with representative homologous prenyltransferases.**

**a,** Structures of monomeric ChlG (top) and other prenyltransferases with PDB codes indicated. ChlG α-helices are numbered. In ChlG, $Mg^{2+}$ ions are placed based on the AlphaFold3 prediction, whereas in UbiA, $Mg^{2+}$ ions from the crystal structure are shown. **b,** Sequence alignment of ChlG and prenyltransferases represented in (**a**) colored by conservation. Substrate entry loops and sequences responsible for ligand binding in ChlG are indicated. Stars indicate amino acid residues involved in the interaction with GGPP in ChlG. DGGGP, digeranylgeranylglycerol phosphate synthase. UbiA-PT, UbiA prenyltransferase domain-containing protein-1.

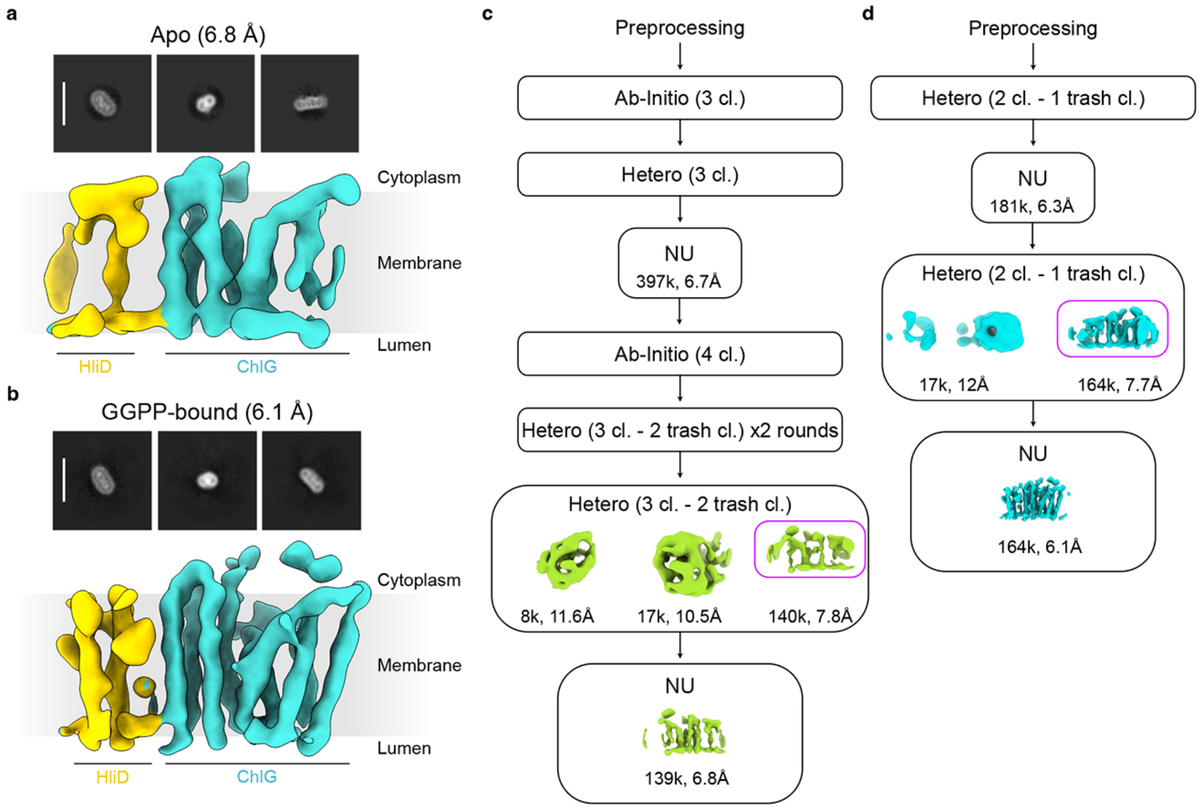


**Extended Data Figure 9. Cryo-EM analysis of the G-D2 complex.**

**a**, Representative 2D classes (top) and cryo-EM map (bottom) of the G-$D_2$ complex from the apo sample. **b**, Representative 2D classes (top) and cryo-EM map (bottom) of the G-$D_2$ complex from the GGPP-bound sample. **c**, Cryo-EM data processing pipeline scheme for the G-$D_2$ complex from the apo sample. **d**, Cryo-EM data processing pipeline scheme for the G-$D_2$ complex from the GGPP-bound sample. For ab initio reconstruction jobs and the final rounds of refinement jobs, the numbers of particles used and the resolutions achieved are indicated in panels **c** and **d**.

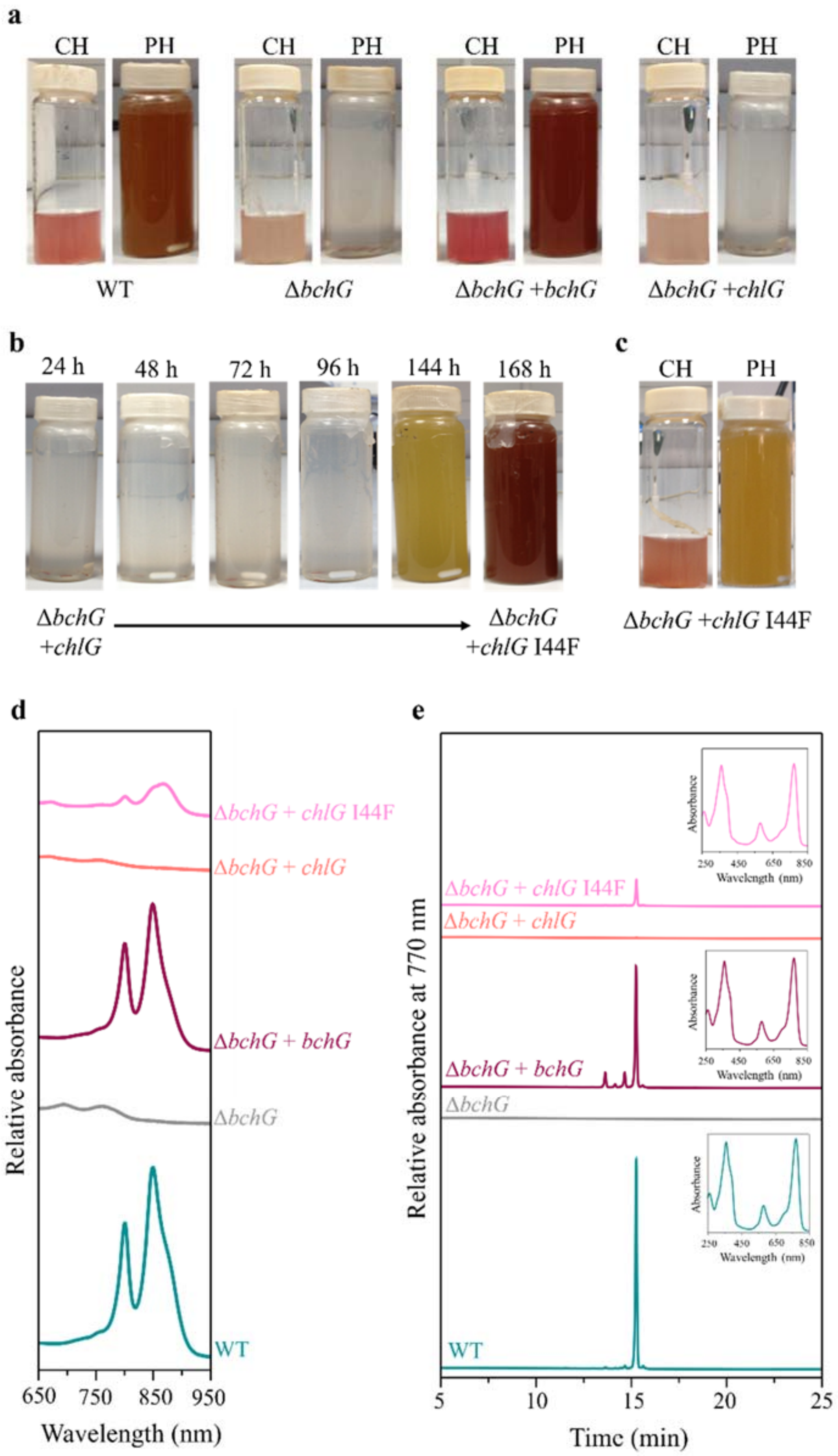


**Extended Data Figure 10. The bacteriochlorophyll-synthesising I44F mutant of ChlG.**

**a**, Cultures of the indicated strains grown under chemoheterotrophic (CH) or photoheterotrophic (PH) conditions. The Δ*bchG*+*chlG* strain grows under CH but not PH conditions. **b**, Adaptive laboratory evolution of the ChlG I44F suppressor mutant. After approximately 6 days incubation under PH conditions, the Δ*bchG*+*chlG* strain started to grow. Sequencing of the *chlG* gene in this strain revealed a single A→T point mutant at position 130 in the gene changing an ATT codon (Ile) to TTT (Phe). **c**, Cultures of the evolved Δ*bchG*+*chlG* I44F strain grown under CH or PH conditions. **d**, Absorption spectra of the strains shown in panels **a** and **c**, showing the lack of RC-LH1 and LH2 complexes in the Δ*bchG* and Δ*bchG* +*chlG* strains. **e**, HPLC quantification of bacteriochlorophyll *a* in the strains shown in panels **a** and **c**. Insets show the spectra of the peak at 15.3 min corresponding to bacteriochlorophyll *a* (where present). The bacteriochlorophyll *a* content of ChlG point mutants analysed in the Δ*bchG*+*chlG* I44F strain described in Table 1 of the main paper was determined in the same way.

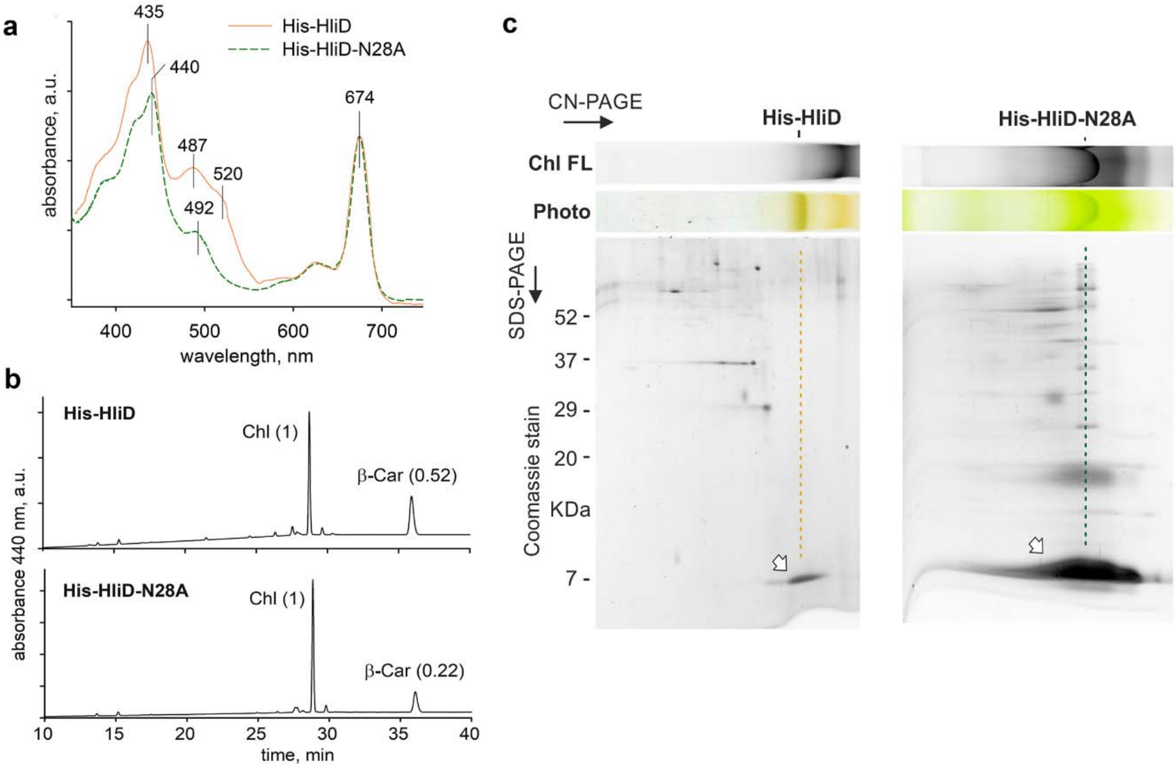


**Extended Data Figure 11. Pigment analysis and 2D-PAGE of the purified HliD and HliD-N28A proteins.**

**a**, Absorbance spectra of His-tagged HliD and HliD-N28A proteins produced in the *Synechocystis* Δ*hliC* genetic background and isolated by immobilised metal affinity chromatography using as Ni-NTA column. **b**, HPLC analysis of isolated protein samples; the relative amount of β-carotene (β-Car) was calculated per 4 chlorophyll (Chl) molecules (values in parentheses). **c**, HliD and HliD-N28A pulldowns, loaded for a similar concentration of Chl (~ 1 µg), were separated by CN-PAGE and then in the second dimension by SDS-PAGE. The resulting gel was stained with Coomassie Brilliant Blue. HliD and HliD-N28A proteins indicated by arrows on the stained gels were assigned according to [1].

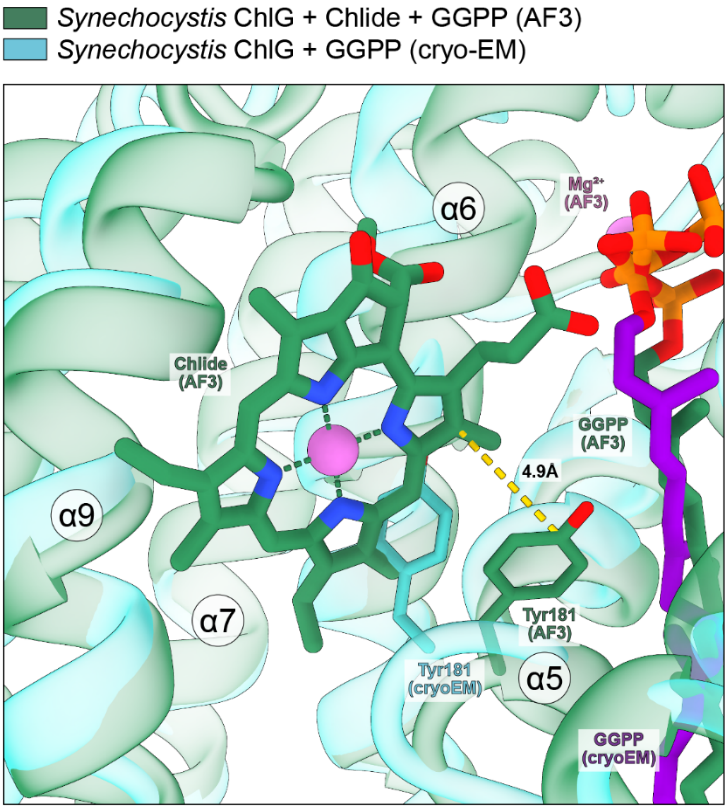


**Extended Data Figure 12. Conformational rearrangement of Tyr181 upon Chlide binding.**

Close-up view of bound Chlide (dark green) in the AlphaFold3 (AF3) model of ChlG (semi-transparent dark green) overlaid with the GGPP-bound ChlG structure determined by cryo-EM (semi-transparent light blue) (also see Figure 3a,b). GGPP (AF prediction, dark green; cryo-EM structure, purple) and $Mg^{2+}$ ions (magenta) are also shown. Tyr181, essential for catalysis, and Ile182, potentially important for substrate selectivity, are shown in stick representation in dark green (AF3) or light blue (cryo-EM). The distance between Tyr181 and the C17-C18 single bond of Chlide is indicated for the AF3 model.

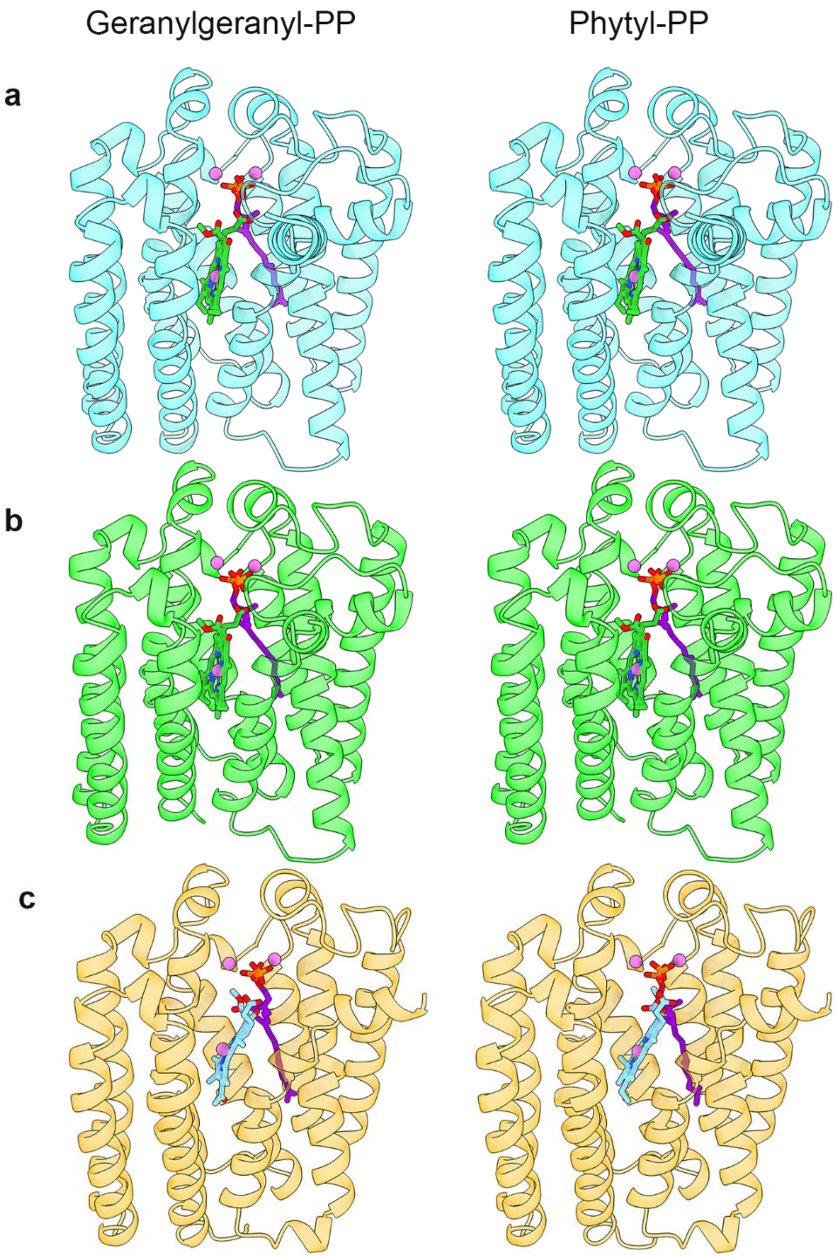


**Extended Data Figure 13. AlphaFold3 (AF3) models of *Synechocystis* ChlG (a), *Arabidopsis* ChlG (b) and *R. sphaeroides* BchG (c) with Chlide/BChlide and GGPP or phytyl-pyrophosphate.**

Lefthand images show AF3 models with Chlide/BChlide (green/light blue) and GGPP (purple); righthand images show AF3 models with Chlide/BChlide and phytyl-pyrophosphate (purple).

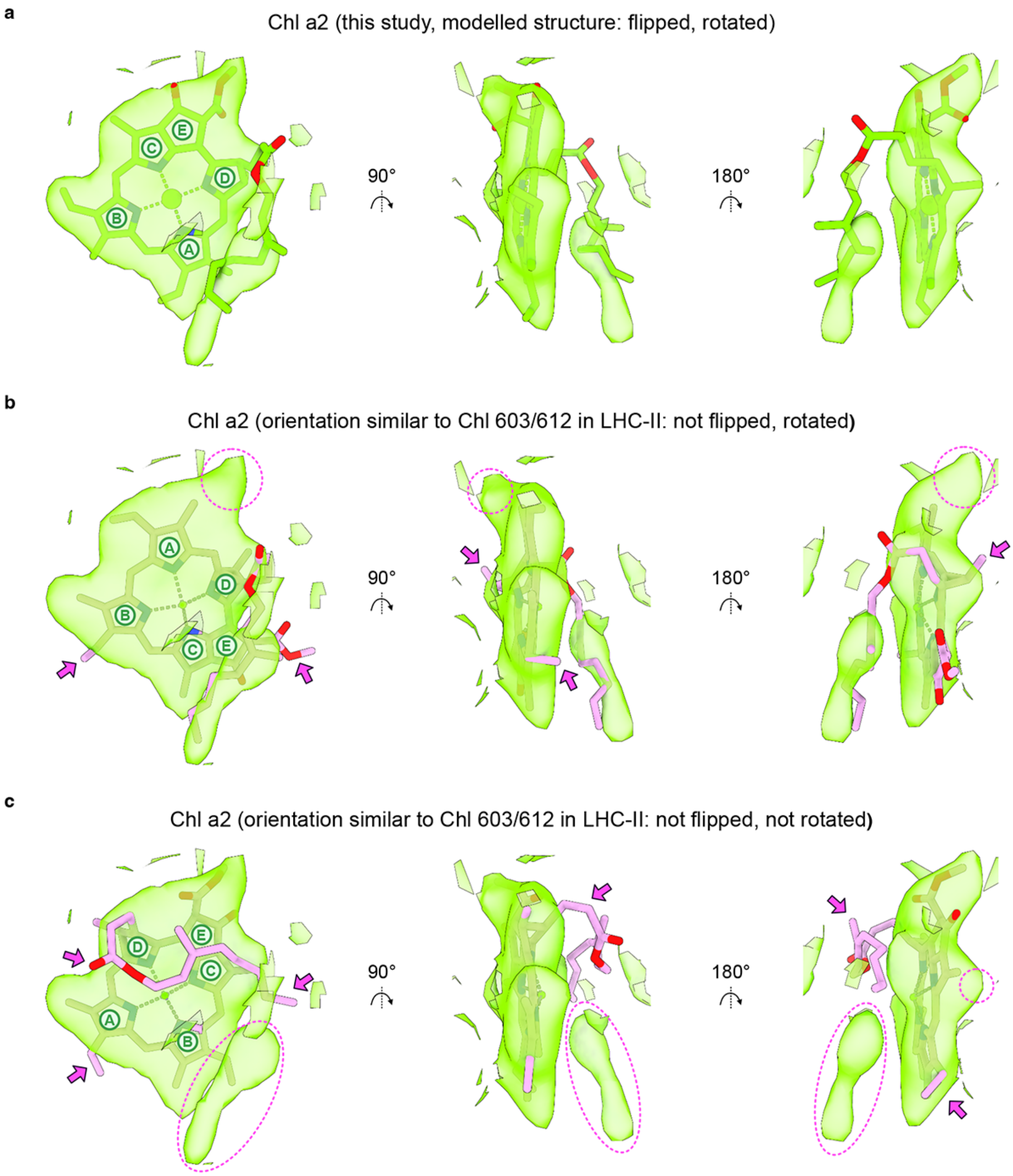


**Extended Data Figure 14. Alternative Chl a2 fittings into the cryo-EM density of the chlorophyll synthase complex.**

**a**, Chl a2 fitted into the cryo-EM density as modelled in this study (PDB:9SAS), macrocycles are indicated. **b** and **c**, Alternative fittings of Chl a2 into the same density as in (**a**) but using the same orientation as in the LHC-II complex (PDB:1RWT). Pink arrows and oval shapes indicate poorly fitted regions of the molecule and empty cryo-EM densities, respectively.

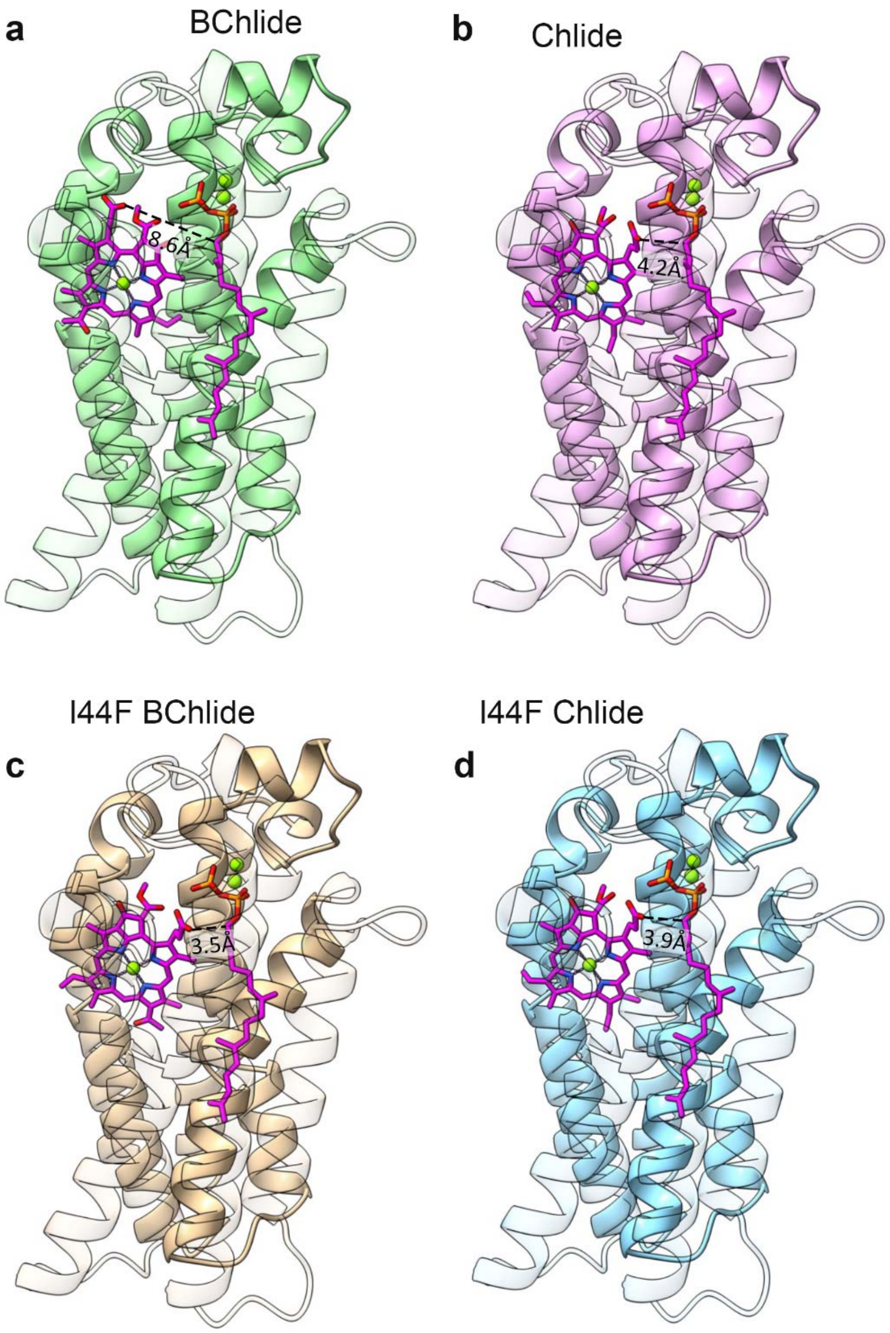


**Extended Data Figure 15. AlphaFold3 (AF3) models of WT ChlG (a,b) and I44F ChlG (c,d) with BChlide (a,c) and Chlide (b,d).**

**a**, AF3 model of WT ChlG with BChlide shows a non-productive binding conformation with the esterifiable carboxyl group 8.6 Å from the terminal GGPP carbon. TMH1-4 are made transparent for clarity. **b**, AF3 model of WT ChlG with Chlide shows the productive binding conformation with the esterifiable carboxyl group 4.2 Å from the terminal GGPP carbon. **c**, AF3 model of I44F ChlG with BChlide shows the productive binding conformation with the esterifiable carboxyl group 3.5 Å from the terminal GGPP carbon. **d**, AF3 model of I44F ChlG with Chlide shows the productive binding conformation with the esterifiable carboxyl group 3.9 Å from the terminal GGPP carbon.

**Table S1. Cryo-EM data collection, refinement and validation statistics (see Methods and Extended Data Fig. 3-5 for details)**

| | ChlG-HliD$_2$-ChlG apo – consensus map (EMDB-54698) | ChlG-HliD$_2$-ChlG apo – consensus map NU2 (EMDB-54696) (PDB 9SAS) | ChlG-HliD$_2$-ChlG GGPP-bound – consensus map (EMDB-54699) | ChlG-HliD$_2$-ChlG GGPP-bound – consensus map NU6 (EMDB-54697) (PDB 9SAU) | HliD$_2$-ChlG apo "half" complex (EMDB-54700) | HliD$_2$-ChlG GGPP-bound "half" complex (EMDB-54701) |
|---|---|---|---|---|---|---|
| **Data collection and processing** | | | | | | |
| Magnification | 165000 | | | | | |
| Voltage (kV) | 200 | | | | | |
| Electron exposure (e–/Å$^2$) | 50 | | | | | |
| Defocus range (μm) | -0.8 to -1.8 | | | | | |
| Pixel size (Å) | 0.68 | | | | | |
| Symmetry imposed | C1 | | | | | |
| Initial particle images after 2D classifications and duplicate removal (no.) | 1258660 | 954046 | 1368937 | 1255494 | 1258660 | 1368937 |
| Final particle images (no.) | 354715 | 210944 | 254925 | 108538 | 139447 | 164515 |
| Map resolution (Å)<br>FSC threshold | 3.47<br>0.143 | 3.38<br>0.143 | 3.7<br>0.143 | 3.49<br>0.143 | 6.78<br>0.143 | 6.06<br>0.143 |
| Map sharpening *B* factor (Å$^2$) | -108 | -65.7 | -149.4 | -52.7 | -544.4 | -610.6 |
| **Refinement** | | | | | | |
| Initial model used | - | AlphaFold; Uniprot IDs: P72932 and Q55145 | - | AlphaFold; Uniprot IDs: P72932 and Q55145 | - | - |
| Model composition<br>Atoms (Hydrogens)<br>Protein residues | - | <br>5874 (0)<br>670 | - | <br>5930 (0)<br>670 | - | - |
| Ligands | - | Myxoxanthophyll: 2<br>Chlorophyll *a*: 4<br>Zeaxanthin: 2<br>Monogalactosyldiacylglycerol: 2<br>Lauryl Maltose Neopentyl Glycol: 2 | - | Myxoxynthophyll: 2<br>Chlorophyll *a*: 4<br>Zeaxanthin: 2<br>Monogalactosyldiacylglycerol: 2<br>Lauryl Maltose Neopentyl Glycol: 2<br>Geranylgeranyl pyrophosphate: 2 | - | - |
| R.m.s. deviations<br>Bond lengths (Å)<br>Bond angles (°) | - | <br>0.002<br>0.662 | - | <br>0.004<br>1.212 | - | - |
| Validation<br>MolProbity score<br>Clashscore<br>Poor rotamers (%) | - | <br>1.53<br>3.05<br>1.67 | - | <br>1.90<br>9.32<br>0.74 | - | - |
| Ramachandran plot<br>Favored (%)<br>Outliers (%) | - | <br>96.07<br>0.00 | - | <br>93.96<br>0.15 | - | - |

Table S1 (continued). Cryo-EM data collection, refinement and validation statistics (see Methods and Extended Data Fig. 3-5 for details)

| | **ChlG-HliD$_2$-ChlG apo – full-map local refinement (EMDB-57787)** | **ChlG-HliD$_2$-ChlG apo – consensus map NU3 (C2 symmetry) (EMDB-57781)** | **ChlG-HliD$_2$-ChlG apo – HliD local refinement map (EMDB-57782)** | **ChlG-HliD$_2$-ChlG apo – ChlG local refinement map (EMDB-57783)** | **ChlG-HliD$_2$-ChlG GGPP-bound – full-map local refinement (EMDB-57788)** | **ChlG-HliD$_2$-ChlG GGPP-bound – consensus map NU7 (C2 symmetry) (EMDB-57784)** | **ChlG-HliD$_2$-ChlG GGPP-bound – HliD local refinement map (EMDB-57785)** | **ChlG-HliD$_2$-ChlG GGPP-bound – ChlG local refinement map (EMDB-57786)** |
|---|---|---|---|---|---|---|---|---|
| **Data collection and processing** | | | | | | | | |
| Magnification | 165000 | | | | | | | |
| Voltage (kV) | 200 | | | | | | | |
| Electron exposure (e–/Å$^2$) | 50 | | | | | | | |
| Defocus range (μm) | -0.8 to -1.8 | | | | | | | |
| Pixel size (Å) | 0.68 | | | | | | | |
| Symmetry imposed | C1 | C2 | C1 | C1 | C1 | C2 | C1 | C1 |
| Initial particle images after 2D classifications and duplicate removal (no.) | 1258660 | 954046 | | | 1368937 | 1255494 | | |
| Final particle images (no.) | 354715 | 210944 | 421888 | 421888 | 254925 | 108412 | 216824 | 216824 |
| Map resolution (Å)<br>FSC threshold | 3.39<br>0.143 | 3.22<br>0.143 | 3.04<br>0.143 | 3.14<br>0.143 | 3.57<br>0.143 | 3.32<br>0.143 | 3.17<br>0.143 | 3.27<br>0.143 |
| Map sharpening *B* factor (Å$^2$) | -108.8 | -77.4 | -110.7 | -123.3 | -134.3 | -70.7 | -118.2 | -111.0 |

**Table S2. iPTM scores for substrates with ChlG from AF3 models**

| Substrate | iPTM to protein |
|---|---|
| GGPP | 0.97 |
| $Mg^{2+}$ | 0.98 |
| $Mg^{2+}$ | 0.92 |
| Chlide *a* | 0.85 |

**Table S3. RMSD of Chlide *a* across the five output AF3 models with respect to best model and distance of carboxyl oxygen from terminal GGPP carbon**

| Model | RMSD (Å) | Distance from carboxyl oxygen to carbon (Å) |
|---|---|---|
| 0 | 0 | 4.3 |
| 1 | 0.659 | 4.8 |
| 2 | 0.261 | 3.3 |
| 3 | 0.467 | 4 |
| 4 | 0.334 | 4.4 |
